# Ligand induced receptor multimerization achieves the specificity enhancement of kinetic proofreading without associated costs

**DOI:** 10.1101/2024.07.26.605371

**Authors:** Duncan Kirby, Anton Zilman

## Abstract

Kinetic proofreading (KPR) is a commonly invoked mechanism for specificity enhancement of receptor signaling. However, specificity enhancement comes at a cost of non-equilibrium energy input and signal attenuation. We show that ligand induced multimeric receptor assembly can enhance receptor specificity to the same degree as KPR, yet without the need for out-of-equilibrium energy expenditure and signal loss. We show how multimeric receptor specificity enhancement arises from the amplification of affinity differences via sequential progression down a free energy landscape. We also show that multimeric receptor ligand recognition is more robust to stochastic fluctuations and molecular noise than KPR receptors. Finally, we show that multimeric receptors perform signaling tasks beyond specificity enhancement like absolute discrimination and aspects of ligand antagonism. Our results suggest that multimeric receptors may serve as a potent mechanism of ligand discrimination comparable to and potentially with more advantages than traditional proofreading.

## I. INTRODUCTION

Cells use secreted ligands and membrane receptors to receive and communicate information about their environment, thereby coordinating cellular activity in response to changing conditions. Ligand-receptor signaling kinetics is the first step in signaling cascades that strongly influences the communication fidelity and the reliability of the resulting cellular responses [1–3]. Many receptor signaling systems require high signaling specificity – the degree to which a cell can respond differently to different ligands based on the differences in their binding affinities to the receptor [3–11].

Receptor specificity is commonly quantified by the ratio of the signaling outputs produced downstream of the receptor by two different ligands, or by the ratio of related quantities such as the average time spent by the receptor in the signaling state or the average occupancy probabilities of the signaling state of the receptor [4, 12– 15]. Another common metric of ligand discrimination is obtained by comparison of the signaling outputs for different ligands to a set threshold that only the agonist ligands cross and activate a cellular response [5, 11, 16, 17]. A paradigmatic mechanism commonly invoked to explain high specificity of many signaling pathways is kinetic proofreading (KPR) [4, 15, 18–21], which has been proposed to explain observations of high specificity for the T cell receptor, FC*ϵ*RI system, RecA assembly, and other systems [15, 18, 19, 21–31]. The KPR receptor mechanism (see Fig. 1A) comprises a sequence of proofreading states that the ligand-bound receptor must pass through sequentially before reaching the signaling state [4]. Each proofreading state creates an opportunity for weakly binding ligands to fall off the receptor before reaching the signaling state, enhancing the difference in the signaling state occupancy between ligands of different binding strengths, and thereby increasing specificity relative to a non-proofreading receptor.

**FIG. 1.**
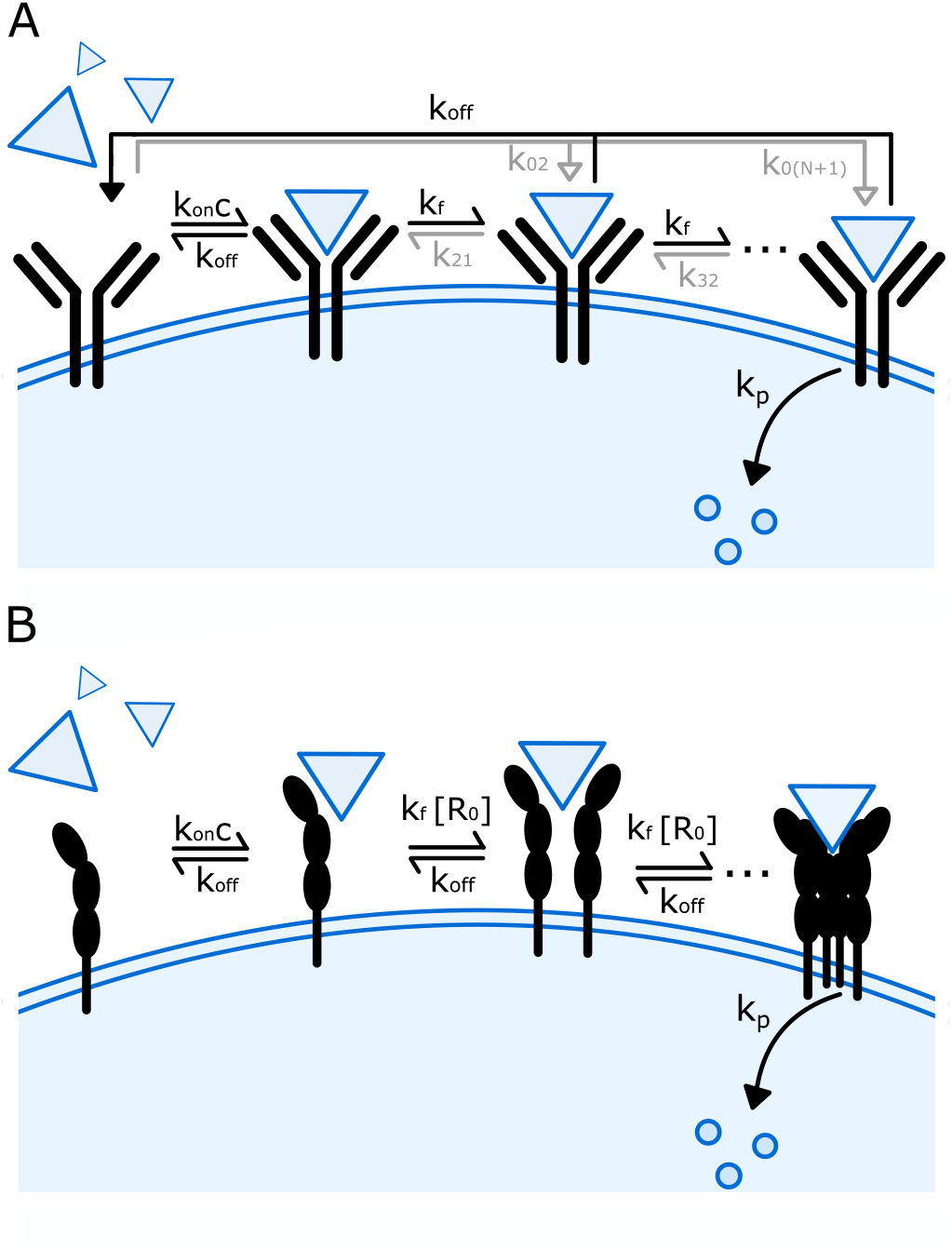
KPR versus multimeric receptors. A) Kinetic proofreading mechanism. The transition rates from the unbound state to all but the first bound state, 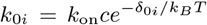, are suppressed by a factor 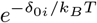, for all *i >* 1 (gray arrows). The transitions *i* + 1 → *i* for all *i >* 0 also proceed at non-equilibrium rate 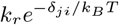 (gray half-arrows). The signaling state of the receptor produces downstream output molecules at rate *k*_*p*_. B) The multimeric assembly mechanism. A multimeric receptor is activated by a ligand crosslinking *N* + 1 monomeric receptor subunits. Receptor subunits are incorporated into the complex at rate *k*_*f*_ [*R*_0_] where [*R*_0_] is the concentration of free receptor subunits on the cell surface.

At equilibrium, where ligand unbinding from any receptor state entails the possibility of a reverse reaction where the ligand binds the receptor, the specificity is limited by the Boltzmann factor ratio of the binding energies of the different ligands [30, 32]. To enhance the specificity beyond equilibrium in KPR, the direct ligand binding to all but the first state in the proofreading chain is suppressed through out-of-equilibrium mechanisms. This typically done through ATP hydrolysis or other biological energy sources, thus preventing short-circuiting the proofreading process [4, 20, 30, 33, 34]. For modeling purposes, the transitions between proofreading states are often assumed to be completely irreversible, with transitions backwards along the proofreading chain fully suppressed (see Fig. 1A) [4, 34].

In addition to the energetic operating costs, other factors constrain the effectiveness of the KPR mechanism. Proofreading works by attenuating the receptor output, which amplifies the relative importance of the intrinsic reaction noise in the output and thereby limits the reliability of downstream ligand discrimination and cellular decision making [4, 35, 36]. Other factors, which limit the utility of KPR in many cellular signaling applications, include the slower response rate, the problem of receptor crosstalk, and extrinsic stochastic fluctuations in the ligand and receptor concentrations [4, 9, 35–39].

In this paper we show that another very common signaling motif, the multimeric receptor, provides receptor specificity enhancement comparable to that of the KPR mechanism. In multimeric receptor signaling, receptor subunits are bound by a multivalent ligand to form an active multimeric signaling complex (see Fig. 1B) [17, 26, 40, 41]. Multimeric receptors are ubiquitous in cytokine and chemokine signaling, as well as GPCR signaling transcription factor interaction networks, and have been implicated in BCR signaling [3, 5, 6, 42–50]. As an example of specificity in multimeric receptor signaling, different Type I Interferons have less than one order of magnitude difference in binding strength to their corresponding dimeric receptor subunits but exhibit two orders of magnitude difference in their potency, and activate different sets of genes [51]. The multimeric Interferon receptor architecture has been identified as crucial to explaining the specificity of these responses [5, 51, 52]. In this paper we show that multimeric receptors can also provide specificity enhancement and ligand discrimination to an extent comparable to the KPR mechanism. Enhanced receptor specificity alone is often not sufficient for reliable ligand discrimination, and we also show that both KPR and multimeric receptors can exhibit variants of “absolute discrimination” and signaling antagonism – signal processing capacities that are important for different cellular decision-making processes. In absolute discrimination, different ligands can be identified independently of their concentration [5, 11, 16, 17]. Antagonism enables ligands to suppress receptor signaling activity, allowing integration of different environmental signals [53, 54].

In contrast to KPR, multimeric receptors do not require non-equilibrium energy input to achieve enhanced specificity relative to a monomeric receptor. We use the framework of non-equilibrium thermodynamics to analyse how the equilibrium and non-equilibrium receptor mechanisms are able to achieve a similar degree of specificity enhancement and absolute discrimination [55].

Our analysis demonstrates that the fundamentally different energetic landscapes of the two mechanisms underlie the functional differences in their signaling performance. Due to these differences, the high specificity of multimeric receptors does not entail non-equilibrium signal attenuation, unlike for KPR [36]. These differences also lead to differences in the sensitivity of each mechanism to molecular noise [1, 8, 37, 56–60], with the multimeric receptor mechanism significantly more robust to molecular noise in functional regimes, enabling higher sensing accuracy.

## II. RESULTS

### A. Description of the models

#### KPR receptor

Kinetics of the KPR signaling receptor are illustrated in Fig. 1A. An unbound KPR receptor binds ligand molecules at rate *k*_on_*c* where *c* is the ligand concentration and *k*_on_ is the association constant. The receptor may then transition forward and backward between the *N* additional bound states at equilibrium rates *k*_*f*_ and *k*_*r*_ respectively. The ligand can unbind from any of the bound states at ligand-specific rate *k*_off_, which returns the receptor to the unbound state. Here we consider the classical minimal KPR model, where the ligand dissociation rate *k*_off_ is the same from all states, determined by the binding energy of the ligand to the receptor. In more generalized models these rates can differ between different states. Overall, the KPR scheme has *N* + 2 states: the unbound state, *N* proofreading states, and the final bound state that produces downstream signaling molecules with rate *k*_*p*_.

Living cells normally contain multiple copies of receptors. We consider *R*_*T*_ total copies of the KPR receptor, but since all transition rates are independent of other receptors the results are unaffected except in Section II D where the presence of multiple copies of the receptor incr<u>eas</u>es the signal-to-noise ratio by the standard factor of 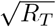 for independent sampling.

#### Multimeric receptor

Kinetics of multimeric receptor assembly is illustrated in Fig. 1B. As mentioned in the Introduction, such multimeric receptor models apply to many signaling systems, including cytokines of the immune system and development [41, 42, 61–64]. Multimerization has also been observed in Fc*ϵ* receptor signaling [25], GPCR signaling [46, 49, 50] as well as BCR signaling [47]. Receptor multimerization can occur differently in other systems, such as the *γ*_*c*_ chain of IL-2 family cytokine receptors [65], where the ligand binds one receptor subunit and the subsequent binding of other receptors to the complex does not create additional ligand-receptor interactions. This alternate form of multimerization may not provide ligand-mediated specificity enhancement.

For simplicity, in figure Fig. 1 we consider a homomultimeric receptor where all receptor subunits are identical. We consider a heteromultimeric receptor in Section II B 1 and Figure 3. The generalized heteromultimeric model is applicable to a wider array of receptors including several cytokine families [5, 61, 64]. The main results of the paper persist for the heteromultimeric model.

In this multimeric receptor model, ligand molecules bind free receptor subunits at rate *k*_on_*c* and unbind at rate *k*_off_ similar to the KPR model. Additional receptor subunits can subsequently associate with the ligand-monomer complex at a rate *k*_*f*_ [*R*_0_], where [*R*_0_] is the in-membrane density of the free receptor subunits with the in-membrane association rate *k*_*f*_, and dissociate from it at the ligand-dependent rate *k*_off_. Unlike for the KPR receptor, the bound receptor complexes are not independent and are coupled through their dependence on [*R*_0_] subject to the conservation of the total number of subunits. We assume that, similarly to *k*_on_ of the threedimensional ligand binding, the two dimensional association rate *k*_*f*_ is primarily determined by the in-membrane diffusion of receptor subunits. As a first approximation, we assume that the in-membrane dissociation rate of receptor subunits from the multimer complexes is equal to the three-dimensional dissociation rate of the ligand from a single subunit, because they are both dominated by breaking the bonds between the ligand and a receptor subunit [5]. This assumption is supported by structural, imaging and systems studies [5, 43, 62]. However, generally this may not be the case, and in Section II B 1 we relax this assumption to allow different in-membrane and three-dimensional dissociation rates. The main results of our paper are independent of this assumption.

A full multimeric receptor composed of *N* + 1 monomers bound to a ligand produces downstream signaling molecules with rate *k*_*p*_; *N* = 0 corresponds to a monomeric receptor which signals immediately upon ligand binding. We reuse the variable *N* here because, as we will show below, both the KPR and multimer mechanisms enhance specificity as *N* increases.

In order to study the thermodynamic forces driving each receptor model, we incorporate non-equilibrium energy inputs in a thermodynamically consistent manner. For general microscopic thermodynamic consistency, the state transition rates must satisfy the following relation [55]:

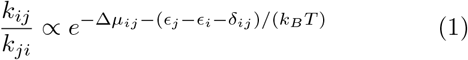

where *ϵ*_*i*_ is the equilibrium energy of state *i, δ*_*ij*_ is the non-equilibrium energy input associated with the transition from state *i* to state *j*, Δ*µ*_*ij*_ is the chemical potential change of this transition, *T* is the temperature and *k*_*B*_ is the Boltzmann constant [55]. This ensures microscopic reversibility of all state transitions, so that for any pair of transition rates (*k*_*ij*_, *k*_*ji*_), between states *i* and *j, k*_*ij*_ ≠ 0 and *k*_*ji*_ ≠0. If *δ*_*ij*_ = 0 for all transitions, the system achieves an equilibrium Boltzmann distribution with the probability of state occupancy 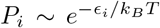 [55, 66–68]. For the KPR receptor, the indices *i* and *j* enumerate the states of the receptor, as illustrated in Fig. 1A. For the multimer receptor the indices *i* and *j* enumerate the states in the multi-dimensional state space defined by the numbers of free and partially multimerized receptor subunits *R*_0_, *R*_1_, …, *R*_*N*+1_. We assume that the receptors are in equilibrium with the bath of ligands with the concentration *c* buffered at a constant value. For the multimer receptor, the change in the chemical potential in Eq. 1 includes both the change in the chemical potential of the ligand solution upon the ligand binding to the receptor, and the change in receptor subunit concentrations upon binding or unbinding of a subunit to or from a multi-subunit complex. Transitions from the unbound state (state 0) to states other than state 1 are prohibited for the multimer because we assume all reactions are bimolecular.

The condition of Eq. 1 leaves undefined the exact dependence of the ligand-receptor binding and unbinding rates on the equilibrium binding energy *ϵ*. In this paper, we use a common choice where the association rates are limited by the diffusion rate of the reactants to the receptor, and the dissociation rates contain the dependence on the difference in the binding energies between states [55, 69]. For the KPR model, this choice results in

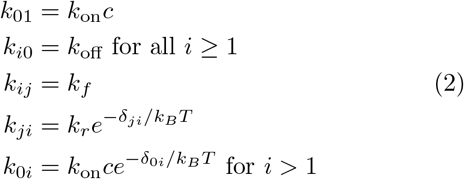

where *k*_*ij*_ is the transition rate of from state *i* to state *j, j* = *i* + 1, and 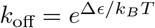 contains the ligand binding energy difference. Transitions from the empty receptor to all but the first bound state in Fig. 1A are suppressed by setting *δ*_0*i*_ ≫ *k*_*B*_*T* for *i >* 1. Suppressing these reactions is essential for specificity enhancement by proofreading [4, 34]. Furthermore, transitions backwards along the chain of bound states are also usually suppressed by setting *δ*_*ji*_≫ *k*_*B*_*T* for *i >* 1 although this is not strictly necessary for specificity enhancement [30, 70].

For the homomultimeric receptor, the transition rates are chosen to be:

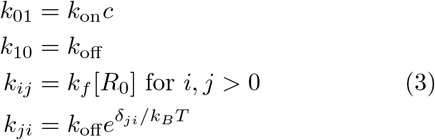

where [*R*_0_] is the density of free receptor subunits on the cell surface. Typical biological values for the rate constants for both receptor models are provided in the S.I.

By analogy with KPR, we include a term *δ*_*ji*_ which alters the transition rate from state *i* + 1 to state *i* for *i >* 1 and represents non-equilibrium energy input relative to the state energies. On a mathematical note, in kinetic schemes without “loops” – such as the homomultimer – the *δ*_*ij*_’s mathematically can be absorbed into the definitions of the state energies; this is not the case anymore for heteromultimers discussed in Section II B 1.

### B. Multimeric receptors can achieve the same specificity as KPR

We consider the classic signal processing task of distinguishing between two ligands which are uniquely identified by their respective receptor unbinding rates *k*_off,1_ and *k*_off,2_ [4]. A common metric of specificity is the ratio of receptor outputs to two different ligands with different unbinding rates *k*_off,1_ and *k*_off,2_ [4, 12, 34]:

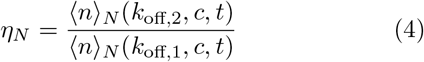

where *n*(*k*_off_, *c, t*) is the total number of output molecules produced by time *t* in response to a ligand with the dissociation rate *k*_off_ at concentration *c*, and⟨·⟩ denotes an average over stochastic realizations of the signaling trajectory. Without loss of generality we set *k*_off,1_ to be the unbinding rate of the weaker binding ligand so that *k*_off,1_ *> k*_off,2_ and the ratio ζ = *k*_off,2_*/k*_off,1_ ≤ 1. With these notations, *η* ≥ 1 and a larger *η* indicates greater specificity. We use ζ = 0.1 throughout this paper as a biologically realistic choice [21, 31, 73, 74].

The discrimination metric defined by Equation 4 implicitly assumes that the two ligands to be distinguished are present at equal concentrations. This is a reasonable assumption in, for instance, some cytokine signaling contexts [5, 52]. In other contexts, the cell may face an additional challenge that the weakly binding ligands are present at significantly higher concentrations than the strongly binding ligands. At low concentrations, where the signaling output is linearly proportional to the ligand concentration, this does not change the utility of the *η*_*N*_ metric, which can be re-normalized by the appropriate ratio of the ligand concentrations.

Another useful metric uses a comparison of the signaling output to a threshold necessary for cellular activation, whereby weak ligands do not cross the threshold [11, 14]. We consider this form of ligand discrimination in Section II C and Fig. 4B in the context of “absolute discrimination”.

For an *N* -step KPR receptor in the irreversible limit (i.e., *δ*_*ji*_ and *δ*_0*i*_ ≫ *k*_*B*_*T*), the mean receptor output is [4]:

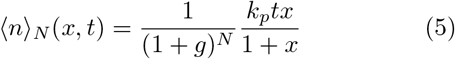

where *x* = *c/K*_*D*_ gives the ligand concentration relative to its dissociation constant *K*_*D*_ = *k*_off_*/k*_on_, and *g* = *k*_off_*/k*_*f*_ measures the strength of proofreading. The specificity enhancement is quantified by the ratio of the specificity for an *N* -step receptor to that of a non-proofreading (*N* = 0) receptor. As established previously, for the KPR receptor in the irreversible limit, the specificity enhancement is [4]:

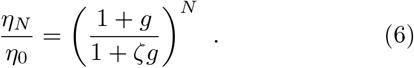

In the strong proofreading regime (*g* ≫ 1) this equation yields the well-known result that the specificity enhancement increases as a power of *N*.

For the multimeric receptors one can similarly compute the specificity ratio as defined in Eq. 4. In the case of a dimeric receptor (*N* = 1), the mean receptor output is:

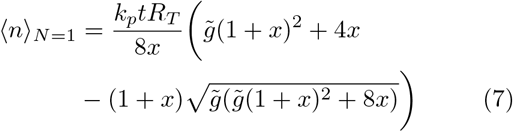

where *x* = *c/K*_*D*_, *K*_*D*_ = *k*_off_*/k*_on_, *R*_*T*_ is the total number of receptor subunits, and *c* is the concentration of the ligand. The parameter 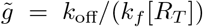, where [*R*_*T*_] is the surface density of all receptor subunits, controls the discrimination strength and is analogous to the proofreading strength *g* = *k*_off_*/k*_*f*_ of the KPR receptor. Both parameters reflect the chance of ligand unbinding before reaching the signaling state, responsible for the specificity enhancement, with larger values indicating a stronger bias away from the signaling state.

The results are summarized in Fig. 2A which shows the specificity as a function of *k*_off_ for the multimeric receptor model and the KPR receptor model in the irreversible limit for *N* ∈ { 0, 1, 2}. Fig. 2A illustrates the central result of this paper: the multimeric receptor is capable of achieving the same maximum specificity as the KPR receptor. At very low values of *k*_off_ (i.e., low *g*) neither model can distinguish between different ligands because in this regime the receptor signaling state is reached quickly (relative to the unbinding time) and it is the diffusion of the ligand to the receptor rather than the unbinding rate that is the limiting factor for signal production. In this limit, the receptor output is essentially independent of *k*_off_ and the ligands are indistinguishable. For high values of *k*_off_ (high *g*), the specificity enhancement relative to the *N* = 0 case reaches the limiting value *η*_*N*_ ≃ ζ^*−N*^ for both receptor models as shown in Fig. 2B.

**FIG. 2.**
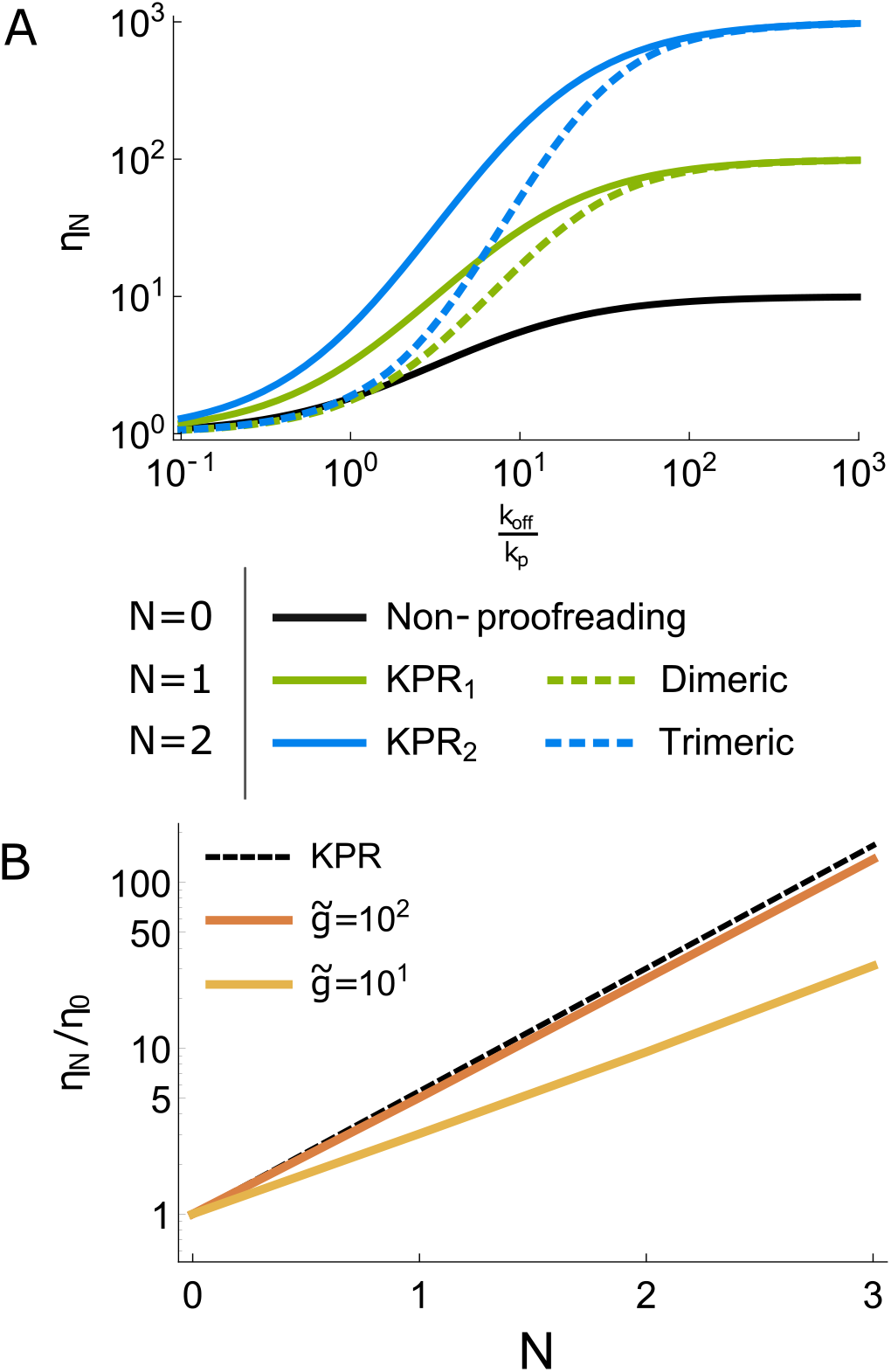
Multimer receptors can yield specificity enhancement of the same magnitude as KPR receptors. A) Receptor specificity for the KPR and multimer models with *N* ∈ {0, 1, 2} as a function of ligand unbinding rate, *k*_off_ (measured in units of *k*_*p*_). Other parameters: *k*_*f*_ [*R*_*T*_]*/k*_*p*_ = 1 for the multimer model, *k*_*f*_ */k*_*p*_ = 1 for the KPR model, *k*_on_*c/k*_*p*_ = 1, and ζ = 0.1. B) The specificity enhancement relative to a monomeric non-proofreading (*N* = 0) receptor, as a function of *N* and 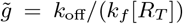. For the multimeric receptor, different choices for the density of receptor subunits, [*R*_*T*_], changes the specificity enhancement. The KPR model is provided for comparison (black dashed line and *g* = 10). Parameters are *k*_on_*c/k*_off_ = 10^*−*1^, *k*_*p*_*t* = 100, and ζ = 0.1. Values of 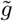 align with those used in [5, 71, 72].

A characteristic feature of the KPR model is that the specificity can be arbitrarily increased by increasing *N* because of the geometric scaling of *η*_*N*_ with *N*, as shown in the black dashed line in Fig. 2B. By contrast, for multimeric receptors, although the specificity enhancement also increases with *N*, the maximum specificity enhancement *η*_*N*_≃ ζ^*−N*^ also depends on the density of the receptor subunits on the cell surface, [*R*_*T*_], as follows from Eq. 7 and is shown in orange and yellow lines in Fig. 2B. The highest specificity enhancement for the multimeric receptor occurs at relatively low [*R*_*T*_], where it approaches that of the the KPR model. For sufficiently large [*R*_*T*_], multimeric receptors do not significantly enhance specificity because signaling complexes form very quickly relative to the characteristic ligand binding time (i.e., 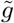 is small, analogous to the weak proofreading regime of KPR) and thus the benefit of multimerization over a monomeric receptor is lost (see also Supplementary Fig. S2). Receptor subunit expression on the cell surface can thus tune the specificity.

Although the values of 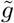 can vary substantially between different systems, typically for cell signaling receptors *R*_*T*_ ∼100 − 1000, although inter-cellular distributions can have long tails; *k*_off_ ∼ 0.001 − 1 *sec*^*−*1^; and *k*_*f*_ ∼ 0.1*µm*^2^*/*(molecules · *sec*) yielding 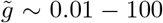 [5, 71, 72]. As an example, the Type I Interferon receptor – a heterodimer – operates at approximately 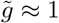 to 10, with a substantial specificity enhancement (see Table S1). The heteromultimeric receptor is also more robust to increasing [*R*_*T*_] and yields specificity enhancement at greater [*R*_*T*_] than the homomultimeric receptor, as discussed in Fig.II B 1.

#### 1. Heteromultimeric receptors

Thus far, we have assumed that the ligand unbinding rate is the same for all subunits, and the in-membrane dissociation rate is equal to the 3D dissociation rate. In this section we relax these assumptions and consider the general case with *N* + 1 distinguishable receptor subunit types that each bind the ligand with a different unbinding rate *k*_off,*α*_ for *α* ∈ {1, *N* + 1}. A common molecular realization of this scenario is when each receptor subunit binds to a distinct part of the ligand (e.g. [51]). Heterodimer kinetics are illustrated in Fig. 3A; models with more than two types of receptor subunits have increasingly more complex kinetic schemes for the formation of the signaling complex. Note the presence of a closed loop in the kinetic scheme of the heterodimeric receptor; this feature persists for heteromultimers with higher *N*.

**FIG. 3.**
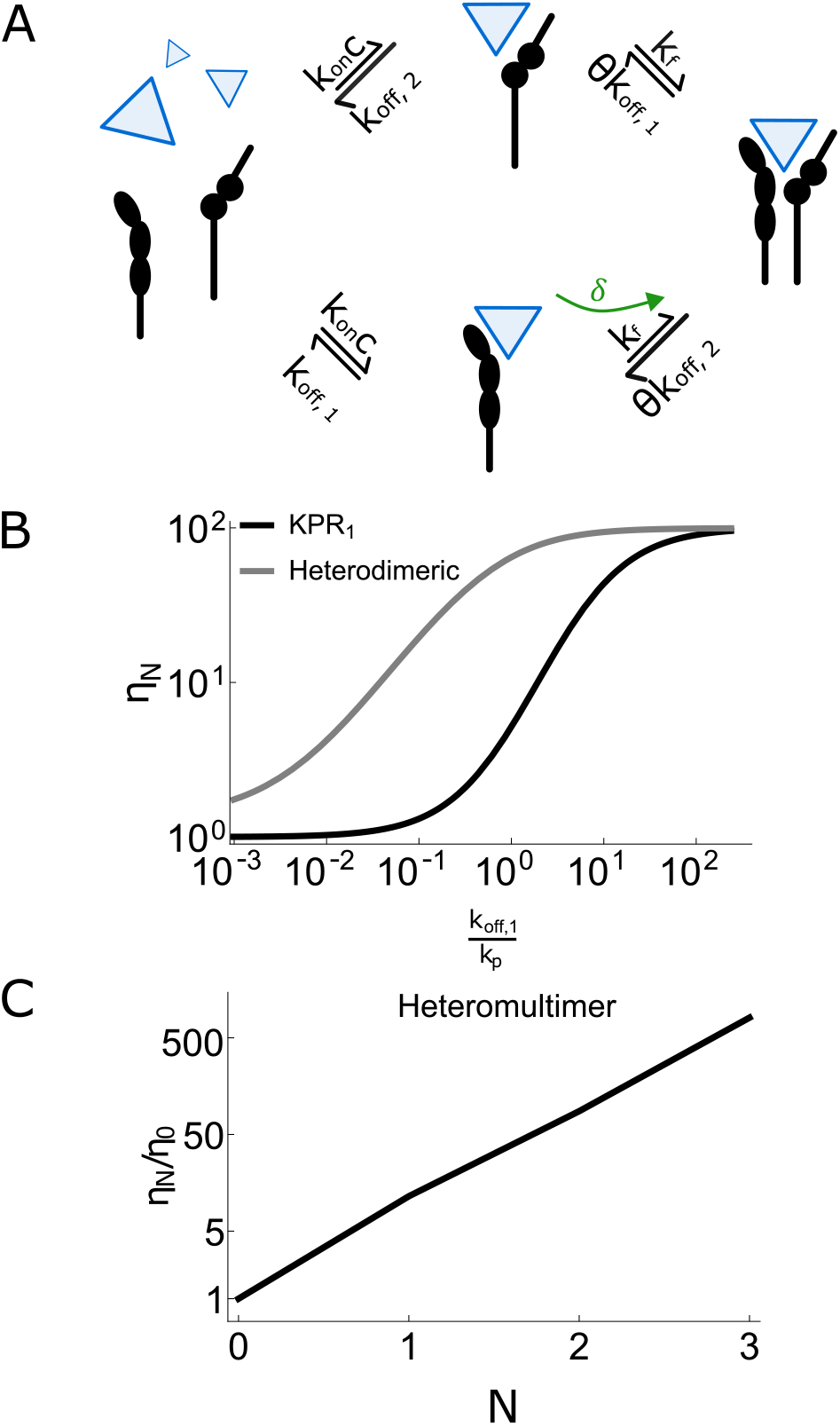
Heteromultimeric receptors exhibit specificity enhancement comparable to KPR. A) The heterodimeric receptor as a prototypical example of the heteromultimeric receptor model. Each receptor subunit has a unique binding affinity for the ligand. The dissociation of a receptor subunit from a complex involving other receptors occurs at in-membrane rate *θk*_off_, different from the three-dimensional ligand-receptor dissociation rate, *k*_off_; *δ*indicates non-equlibrium energy input. See S.I. for the detailed mathematical description of the model. B) The heterodimeric receptor can achieve the same sensitivity enhancement as the 1-step KPR model, even at higher receptor numbers. Parameters: *k*_*f*_ [*R*_*T*_]*/k*_*p*_ = 10, *k*_off,2_*/k*_*p*_ = 2, *θ* = 0.5, *δ*= 0; all other parameters the same as Fig. 2A. C) The heteromultimeric receptor specificity enhancement, *η*_*N*_ */η*_0_, scales similarly to the KPR and homomultimeric models shown in Fig. 2B. Parameters: *k*_*f*_ [*R*_*T*_]*/k*_*p*_ = 1, 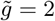 for the first type of receptor subunit (i.e., *N* = 0),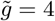 for the second type of receptor subunit (when *N* = 1), 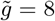 for the third type of receptor subunit, 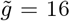 for the fourth type of receptor subunit, *θ* = 0.5, *k*_on_*c/k*_*p*_ = 0.02, and simulations run to steady state. For the heterodimer the values of 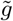 where specificity enhancement is observed can be lower than for the homodimer.

As shown in Figure 3B and C, the heteromultimeric receptor model acts similarly to a homomultimeric receptor or KPR receptor, demonstrating the same degree of specificity enhancement and exponential scaling of *η*_*N*_ with *N*. Moreover, the heterodimeric receptor maintains high specificity at higher total receptor density, [*R*_*T*_], than a comparable homodimeric receptor. Finally, the results also hold in the case of different in-membrane versus three-dimensional dissociation rates modeled by parameter *θ* (see Section II A) and are robust in a wide range of values of *θ*.

### C. Absolute discrimination

We have shown that multimeric receptors are capable of the same degree of specificity enhancement as KPR. However, specificity enhancement *per se* is not sufficient for robust signal discrimination in many contexts where different ligands must be distinguished unconfounded by the potential concentration differences [5, 11, 16, 17].

Ligand discrimination independent of ligand concentration – known as “absolute discrimination” – is not possible with a non-proofreading monomeric receptor (*N* = 0) because the receptor output saturates to the same level dictated by the total number of receptors, *R*_*T*_, independently of *k*_off_; thus the weak binding can be compensated by higher concentration [5, 11, 16, 17].

The concept of absolute discrimination was originally introduced in [11, 16] in order to explain the selfnonself discrimination by TCRs. These works introduced kinase-mediated negative feedback to the KPR scheme, resulting in the “adaptive sorting” mechanism. The notion of absolute discrimination was later adopted in the context of dimeric receptors to explain their ability to discriminate between different ligands – such as different Type I Interferons – that act through the same receptor but elicit different cellular outcomes [5, 17].

A sufficient condition for absolute discrimination is that the maximum receptor output depends on *k*_off_ in some concentration range [5, 16, 17]. In this case, a response threshold can be chosen such that the weaker binding ligand of a pair of ligands can never induce a cellular response at any concentration, thus enabling absolute ligand discrimination, as illustrated in Fig. 4A [11, 16, 17, 75].

**FIG. 4.**
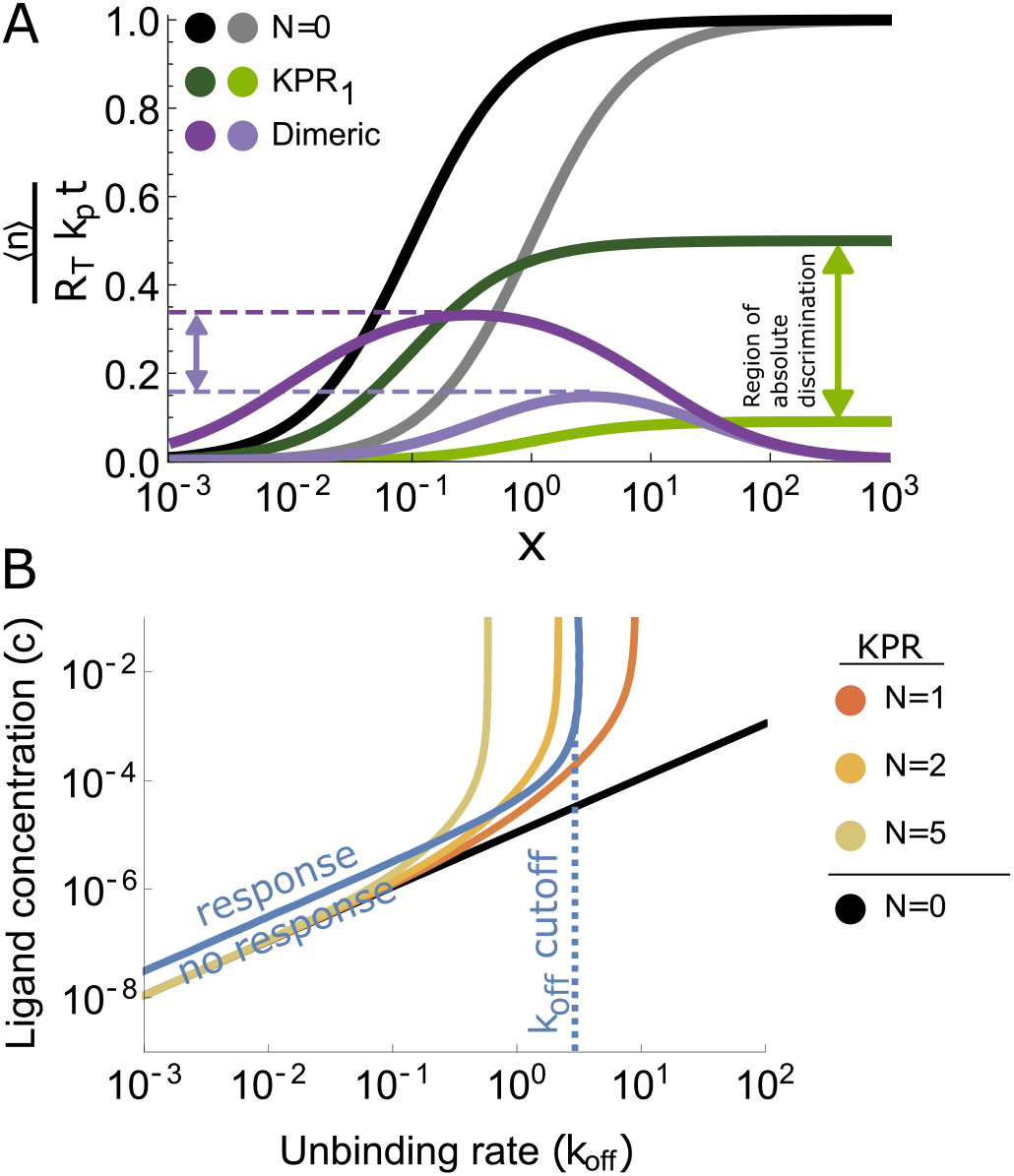
Threshold crossing and absolute discrimination. A) The dose-response curves for the non-proofreading monomeric receptor (*N* = 0, black & gray), 1-step KPR receptor (green), and heterodimeric receptor (purple) in response to two different ligands with 10-fold difference in affinity (ζ = 0.1). The response to the ligand with a larger *k*_off_ is given in the lighter shade for each model. The difference in maximum response between the two ligands allows absolute discrimination in the (*N* = 1) KPR and dimeric receptor models. The region of absolute discrimination for the heterodimer covers four orders of magnitude of ligand concentrations, a greater range of concentrations than an equivalent homodimeric receptor (see S.I). The response ⟨*n*⟩ is normalized by *k*_*p*_*tR*_*T*_. Parameters: *x* = *k*_on_*c/k*_off_, *g* = 10 for the KPR model, 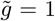 for the first receptor subunit and 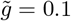 for the second receptor subunit in the dimeric receptor, *k*_off_*/k*_*p*_ = 1 for the *N* = 0 receptor, and *k*_*p*_*t* = 100. B) Ligand concentration *c* (in units of *k*_*p*_*/k*_on_) required to achieve a desired response – fixed at 30 output molecules per receptor – as a function of the ligand unbinding rate *k*_off_ (in units of *k*_*p*_). For the heterodimeric receptor, we fix one of the receptor affinities (*k*_off_) while varying the other receptor affinity. For all models except *N* = 0, there is a maximum *k*_off_ above which the desired response cannot be achieved at any concentration – a hallmark of “absolute discrimination”. For comparison with the adaptive sorting model and the homodimeric receptor, see Fig. S2. Parameters: *k*_*p*_*t* = 300 for all models; *k*_*f*_ */k*_*p*_ = 1 for the KPR model; *k*_*f*_ [*R*_*T*_]*/k*_*p*_ = 2 and *k*_off,1_ is varied while *k*_off,2_*/k*_*p*_ = 10^*−*4^ for the heterodimeric receptor model.

For KPR and its derivative mechanisms, the absolute discrimination region persists to infinite concentrations as illustrated in Fig. 4A. For multimeric receptors, the response to either ligand is shut down at high ligand concentrations due to the competition for free receptor subunits, and absolute discrimination is obtained in a bounded range of concentrations (Fig. 4A). The absolute discrimination range is wider for heteromultimeric receptors due to a broader peak in the dose-response curve (see Fig. 4 and Fig. S2G) [5, 75].

Amplitude based absolute discrimination based on differences in the maximum receptor output has been shown to explain functional differences between Type I Interferons [5, 52] and is consistent with the observed differences in signaling outcomes between engineered diabodies in erythropoeitin receptor systems [41].

For the KPR receptor, the maximum normalized response is max_*c*_[⟨*n*⟩_1_] = *R*_*T*_ *k*_*p*_*t*(1+*g*)^*−N*^ (from Eq. 5). For the multimeric receptor there is no closed form solution for the maximum response for all *N*, but the maximum for the case of *N* = 1 provides an informative example:

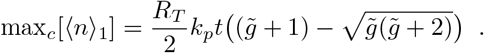

The analagous expression for the heterodimer is provided in the S.I.

Fig. 4B evaluates the absolute discrimination using an alternative metric of discrimination to *η*_*N*_ of Eq. 4 by comparing the signaling output ⟨*n*⟩ to a set threshold. The regime where the signaling output ⟨*n*⟩ is larger than the threshold is considered to generate a response. The unproofread monomer (*N* = 0) does not exhibit any response cutoff at any *k*_off_ value. By contrast, increasing *N* both for KPR and multimer receptor models creates an increasingly sharp cutoff for weak binding ligands, seen as a vertical asymptote in the response contours as *k*_off_ becomes large. For further comparison of the performance with adaptive sorting, see S.I. and Fig. S2.

It is interesting to note that multimeric receptors – because their response shuts down at high concentrations – can also act as a band pass filter by preventing cellular response to high ligand concentrations (Fig. S2F). This may be important for applications like protein engineering for cytokine drug development [76].

Another form of signaling response modulation is ligand antagonism, where the response to a strongly binding ligand (agonist) is diminished when ligands of similar but slightly weaker binding strength are also present (antagonists), allowing one ligand to filter or modulate the response to another ligand [53, 77, 78]. In the S.I. we show that both multimeric and KPR receptors can exhibit antagonism mediated by competitive binding in ligand mixtures. These receptor models may therefore allow cross-regulation between ligands via ligand antagonism, providing another mechanism for complex cellular signal processing. We note that in the important immunological example of the T cell receptor, antagonism appears to be mediated by internal negative feedback through adaptive sorting rather than through competitive ligand binding, and the experimentally observed antagonistic effect is much stronger than is possible to explain with competitive binding via the KPR or multimeric receptor models [11, 56].

### D. Multimeric receptor performance is more resilient to intrinsic noise than KPR

Intrinsic stochasticity of the reaction kinetics in both the KPR and multimer models leads to variance in the receptor output, which can significantly limit the accuracy of downstream signal processing [1, 11, 31, 36, 57, 59, 69, 72, 79]. Such noise affects the precision of ligand identification and classification [10, 36, 69], accuracy of decision making with many possible responses (sometimes referred to as functional plasticity) [3, 5, 43], and the identification and disentanglement of mixtures of ligands presented to the receptor [69, 80–83]. KPR is known to be particularly vulnerable to noise because the specificity enhancement is accompanied by signal attenuation [36, 59, 70]. By contrast, dimerization has been studied as a method of noise reduction [84]; here, we provide a broader analysis for multimerization.

The noisiness of a signal is commonly quantified by the signal-to-noise ratio (SNR):

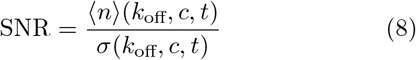

where *n* is the number of output molecules produced by the receptor, *σ* is the standard deviation of this output, ⟨·⟩ denotes the average over stochastic realizations of the signaling dynamics, and both ⟨*n*⟩ and *σ* are computed at time *t* for a ligand at concentration *c* and with the unbinding rate *k*_off_. Derivations of *n* and *σ* for both receptor models are provided in the Methods.

As shown in Fig. 5 the SNR of the KPR receptor monotonically decreases with *N* because the output from the receptor is attenuated by each additional proofreading step, increasing the relative importance of intrinsic fluctuations [4, 36]. By contrast, the SNR of the multimeric receptor model improves as *N* increases, at least in the low concentration regime (i.e. *c < K*_*D*_; additional parameter regimes are shown in Fig. S3). Although the output of the multimer receptor is also attenuated with *N* for 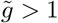 because fewer active signaling complexes can be formed from the same *R*_*T*_ receptor subunits (see Eq. 13), the multimeric signaling complex becomes more energetically stable as *N* increases (see Section II E), which partially compensates for signal attenuation. This suggests that multimeric receptors may operate better at higher *N* while KPR may favour low *N*.

**FIG. 5.**
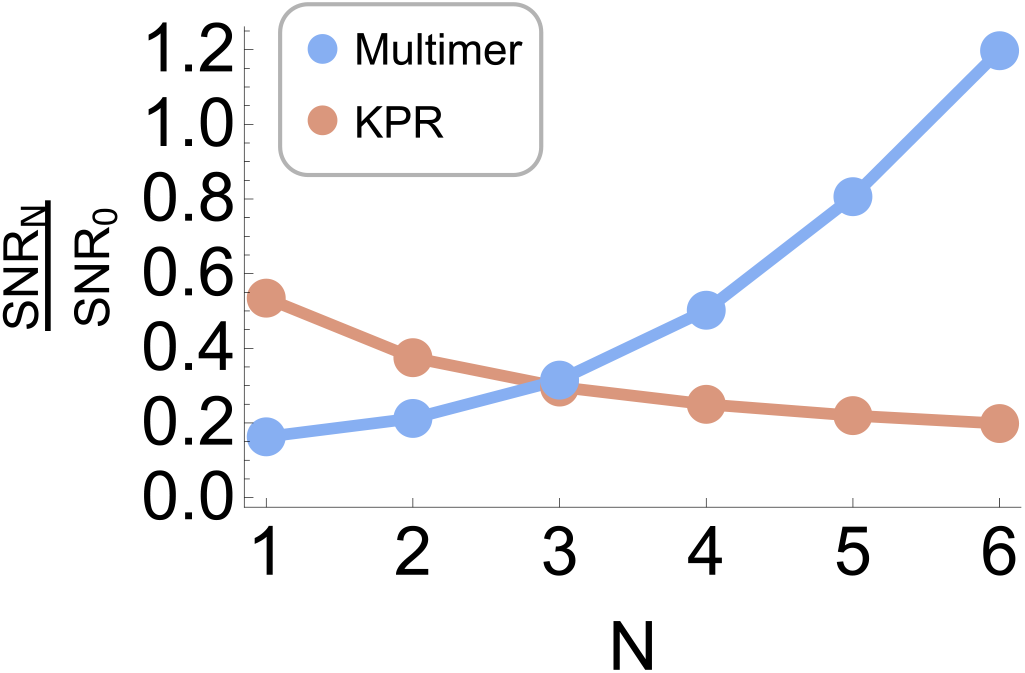
Signal-to-noise ratios for KPR and multimer receptors. The SNR of each receptor model relative to that of the *N* = 0 model, as a function of *N*. Parameters: *R*_*T*_ = 100(*N* + 1) for the multimer so that for all *N* at most 100 signaling complexes can be formed; *R*_*T*_ = 100 for the KPR receptor to match the multimer; other parameters are *k*_on_*c/k*_*p*_ = 10^*−*2^, *k*_*p*_*t* = 100, and ζ = 0.1. For both models, 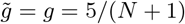 (moderate proofreading strength).

### E. Receptor energy landscape determines specificity

An important rationale for KPR as a specificity enhancement mechanism is that it provides a minimal model for incorporating the non-equilibrium energy input that increases the ratio of the signaling state occupancies between ligands beyond the equilibrium Boltzmann factor, 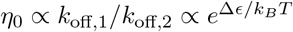, where Δ*ϵ* = *ϵ*_2_ − *ϵ*_1_ is the difference in the binding energy between two ligands [4]. Given that the multimeric receptor assembly occurs under equilibrium conditions, it may seem perplexing that multimeric receptors can achieve specificity greater than the Boltzmann factor. To better understand the underpinnings of this apparent puzzle we examine the energy landscape for each receptor mechanism.

#### KPR receptor

While the KPR receptor is not usually considered at equilibrium, its equilibrium energy landscape provides a contrastive example to the energy landscape of the multimeric receptor and helps to explain the differences in signaling performance between the two. For the standard KPR, where unbinding rates from all bound states are equal to *k*_off_, the energy of the system in each bound state is the same and is equal to

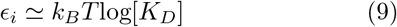

where *K*_*D*_ = *k*_off_*/k*_on_ (See Eq. 19 of the Methods for discussion). Equation 9 expresses the fact that the binding energy of the ligand is the same in all KPR states, with the same dissociation rate *k*_off_, because 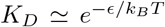 [4]. This is illustrated in Fig. 6A. At equilibrium, the ratio of the occupancies of the signaling state by the two ligands is given by the equilibrium Boltzmann factor 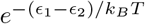. Thus, the KPR receptor must operate out of equilibrium to achieve enhanced specificity beyond the Boltzmann factor via an out-of-equilibrium suppression of the direct ligand binding to any but the first bound state in the proofreading chain.

**FIG. 6.**
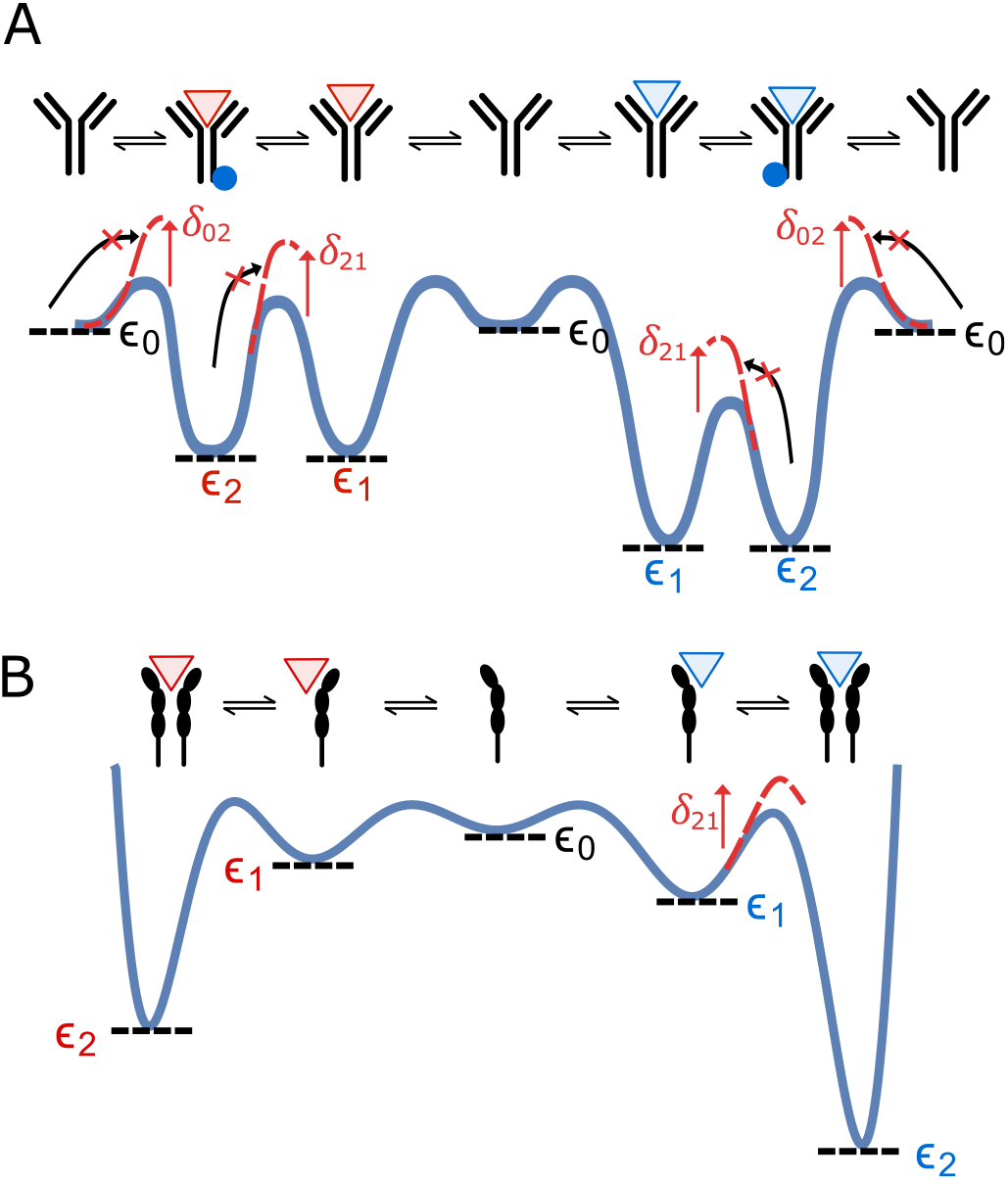
Energy landscape for KPR and multimer receptor signaling mechanisms. A) The *N* = 1 KPR receptor has three possible states (one unbound and two bound states), with the corresponding energies *ϵ*_0_, *ϵ*_2_, *ϵ*_2_ respectively. States 1 and 2 are of equal energy for a given ligand determined by *k*_off_ (see text). Non-equilibrium energy input *δ*suppresses transitions from the unbound state directly into the signaling state (state 2) and also the transition from each bound state to the previous bound state (e.g., state 2 to state 1), generating a non-equilibrium distribution of state occupancy. B) Each receptor subunit of the dimeric receptor model can be in one of the three states in the multimerization chain: unbound (state 0), bound by ligand (state 1), or bound in a dimeric complex (state 2). The stronger ligand (blue) is energetically more stable than the weaker binding ligand in state 1 because of the difference between *ϵ*_1_ for each ligand; for state 2 this difference is even larger. Non-equilibrium driving *δ*_21_ is shown here as an example of how the system can be driven out of equilibrium.

#### Multimeric receptor

By contrast, for the multimeric receptor the energy of a state *i* in the multimerization chain is – in the absence of binding cooperativity – the sum of the binding energies of the ligand to each of the receptor subunits. More precisely, expressed in terms of the kinetic constants, the energy of a receptor in state *i* is:

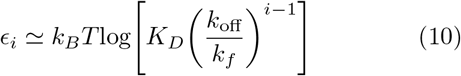

where *K*_*D*_ = *k*_off_*/k*_on_ and *i* is the receptor state in the multimerization chain as described in Section II A. Therefore, for multimeric receptors with *N* + 1 subunits, the signaling states for two different ligands differ in their energy by a factor of *N* + 1 compared to the difference for a non-proofread receptor. This result is illustrated in Fig. 6B, which shows how the stronger binding ligand becomes increasingly energetically favoured over the weaker binding ligand as it binds more receptor subunits. The multimeric receptor therefore discriminates between ligands by amplifying the difference in binding energies between ligands via a progression down the energy landscape. The energy landscape of the multimeric receptor also helps to explain why the multimeric receptor is more robust to stochastic fluctuations: increasing energetic stability of the bound states partially compensates for the signal attenuation as described in Section II D. Although multimeric receptors can achieve enhanced specificity already at equilibrium, it can be further enhanced by appropriate non-equilibrium driving (see Fig. S5 in the S.I.).

## III. DISCUSSION

In this paper we have shown that a multimeric receptor acting via ligand-induced multimerization of receptor subunits can achieve specificity enhancement paralleling that of the paradigmatic kinetic proofreading mechanism (Section II B and Fig. 2). We also showed that multimer receptors are, similar to KPR, capable of performing additional signaling tasks – such as variants of absolute discrimination and antagonism (Section II C), extending previous work [5, 17, 75].

However, the fact that both models can perform analogous functions does not mean that these mechanisms are interchangeable in cell signaling. In particular, multimeric receptors achieve absolute discrimination in a range of concentrations because the response shuts down at high concentration. This is different from KPR and previously discussed “adaptive sorting” models where absolute discrimination is based on the receptor-triggered negative feedback. This feature may provide an additional “knob” for tuning response specificity. The degree of antagonism also differs between the multimer, KPR, and negative feedback-based absolute discrimination mechanisms (as discussed in S.I.).

Another important difference between the KPR and multimer models is that for the multimeric receptor the specificity enhancement explicity depends on the receptor density and is higher at lower receptor density. Experimentally, it has been observed that cells expressing abundant dimeric receptors for certain cytokines may fail to distinguish between cytokines acting through the same receptor [51, 85]. The dependence on *R*_*T*_ demonstrated in Fig. 2 and Fig.3 may thus be important for explaining these experimental observations.

We also showed that the multimeric receptor is more robust to molecular noise. For KPR, the effects of the intrinsic signaling noise become progressively more significant as proofreading steps are added because KPR attenuates the signal. By contrast, the multimeric receptor is increasingly more robust to noise as more receptor subunits become involved because the increased energetic stability of the complex partially compensates for signal attenuation (Fig. 5) [84]. These differences arise from the different energy landscapes of both receptor mechanisms.

Another crucial difference is that, unlike KPR, the multimeric receptor can be highly selective already at equilibrium because the signaling complex becomes more energetically stable with additional bound subunits (Section II E). Thus, multimeric receptors achieve equilibrium energetic specificity enhancement by amplification of the receptor binding strength. By contrast, the equilibrium energy landscape of the KPR receptor is flat, and it achieves kinetic specificity enhancement via nonequilibrium energy input to attenuate the occupancy of the signaling state. In the related problem of proofreading in biosynthesis, specificity enhancement via nonequilibrium modulation of transition rates has been referred to as kinetic discrimination, to distinguish it from discrimination based on an energy landscape which was referred to as energetic discrimination [86]. Multimeric receptors and KPR receptors may represent an example of this difference in cell signaling mechanisms.

Energetic discrimination and kinetic discrimination are not necessarily mutually exclusive. Distributive proofreading is another mechanism in which specificity is enhanced by a combination of irreversible chemical reactions and spatially separated signaling components [87, 88]. Other signal processing mechanisms like information ratchets [89], non-equilibrium allosteric sensing [90], and serial engagement models [91] may also be viewed in this energetic-kinetic framework. In general, the type of signaling schemes considered can be combined in various ways in more complex signaling models [20, 30, 31, 67]. Multimeric receptor specificity can also be further tuned by non-equilibrium energy input (S.I.) but the full analysis of the complete non-equilibrium heteromultimer model lies outside the scope of this work.

Receptor signaling mechanisms have often been viewed as serving a single purpose, such as specificity enhancement by KPR, but we have demonstrated (Section II C) that both KPR and multimeric receptors are multifunctional with overlapping yet non-redundant signal processing capacities. Additionally, both receptor mechanisms are likely capable of producing multiple outputs from different receptor bound states, enabling combinatorial signal processing [69, 80, 81]. We expect that a deeper theoretical understanding of the range of signal processing functions that can be implemented by a given signaling mechanism will be useful in elucidating different roles of pleiotropic receptors and other receptor signaling mechanisms.

## IV. METHODS

### A. Mean receptor signaling dynamics

The KPR receptor states are enumerated by index *i*, with *i* = 0 representing the unbound state and *i* = *N* + 1 the signaling state. The probability of the receptor to be in state *i* and for *n* output molecules to have accumulated at time *t* is 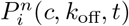. For a single receptor, the dynamics of this probability distribution is described by the Master equation

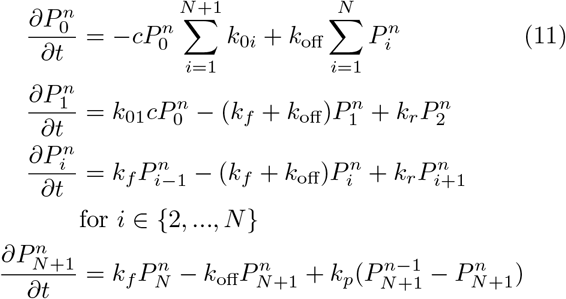

where *k*_0*i*_ are the association rates of the free ligands to a receptor in state *i, k*_*f*_ is the rate at which the receptor transitions from state *i* to state *i* + 1 for *i >* 0, *k*_*p*_ is the rate of signal production from the terminal receptor state; we assume that the ligands are in a large extracellular volume so that *c* remains constant. For the irreversible KPR model *k*_0*i*_ = 0 for *i >* 1, *k*_01_ = *k*_on_, and *k*_*r*_ = 0. The mean and variance of the signal molecule, ⟨*n*⟩ and 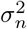, can be obtained from this Master equation by solving the above system of equations for the steadystate receptor occupancy probabilities [36]. For a system of *R*_*T*_ independent receptors, the average number of receptors in state *i* is *P*_*i*_*R*_*T*_. Accordingly, ⟨*n*⟩ and 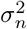, both increase by a factor *R*_*T*_.

The transition rates for the multimeric receptor model depend extensively on the available receptor subunits (e.g., *k*_*i*(*i*+1)_ = *k*_*f*_ [*R*_*i*_][*R*_0_] for *i >* 0, where [*R*_0_] is the concentration of free receptor subunits). This makes it impractical to use a Master equation approach to compute statistics of the signaling dynamics. To compute the mean receptor output we can use equilibrium chemical kinetics because our system involves large numbers of molecules:

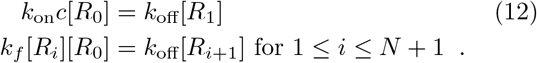

The mean signaling dynamics of the adaptive sorting model are similarly modeled using chemical kinetics (see S.I.). For the case *N* = 1 (a dimeric receptor), the solution to Eq. 12 is given by Eq. 7. The solution for the case *N* = 2 is too cumbersome to provide here but is available in the associated Mathematica notebook (see S.I.). In general, Eq. 12 gives that

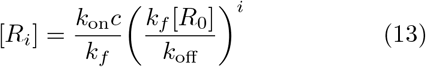

and [*R*_0_] is obtained from the conservation equation:

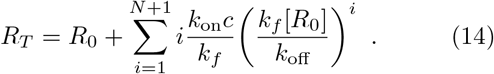

Equation 14 can be solved numerically and reveals that *R*_0_ decreases geometrically with *N* so that we conclude the signaling state [*R*_*N*+1_] is attenuated as *N* increases.

### B. Energy landscapes for the receptor models

A state of a system of of *R*_*T*_ receptors is characterised by a set of variables ***θ*** = {*R*_0_, *R*_1_, …, *R*_*N*_} where *R*_*i*_ denotes the number of receptors in the *i* state in the case of the KPR receptor, or the number of receptor complexes with *i* bound subunits in the case of homomultimer receptor. At thermal equilibrium, the probability of a state ***θ*** is given by the grand canonical distribution

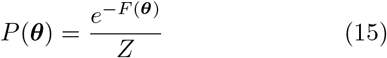

Where 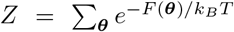 is the partition function. *F* (***θ***) is the grand-canonical free energy functional

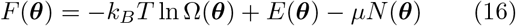

Where 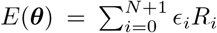 is the energy of the ***θ*** configuration, 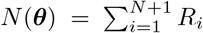 is the number of bound ligands, Ω(***θ***) is the multiplicity, *T* is the temperature, and *µ*≃ *k*_*B*_*T* ln *c* is the chemical potential of the ligands in solution outside the cell.

The entropy, *S* = *k*_*B*_lnΩ, is the logarithm of the number of ways of placing the collection of *R*_*i*_ onto the cell surface of area *A*. In the dilute limit,

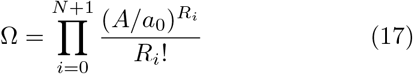

where *a*_0_ is a typical molecular dimension [92– 94]. Applying Stirling’s approximation yields *S* ≃ *k*_*B*_Σ_*i*_*R*_*i*_(ln[*A/*(*R*_*i*_*a*_0_)] + 1).

For large *R*_*T*_ ≫ 1, the probability distribution is dominated by the value ***θ*** = ***θ***^∗^ that minimizes *F* (***θ***),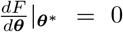 with the constraints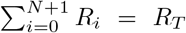 for the KPR receptor and 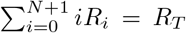 for the multimeric receptor to ensure conservation of the total number of receptor subunits. In the thermodynamic limit *R*_*T*_ → ∞, the thermodynamic grand canonical free energy is *F* (*R*_*T*_, *µ, T*) → *F* (***θ***^∗^) and the 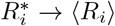, the true ensemble average. For finite *R*_*T*_, 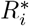 serves as a mean field approximation to the true thermodynamic averages (see also next section). In practice, the thermodynamic expressions are in very good agreement with exact finite size stochastic simulations as shown in the S.I.

Performing the minimization of the free energy for the multimeric receptor by taking the derivative with respect to each *R*_*i*_, we get:

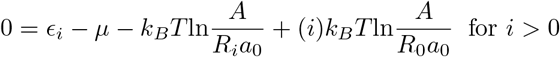

where we have set *ϵ*_0_ = 0 without loss of generality. Comparing these equations to the mean field mass action kinetics, Eqs. 12, we obtain the general formula [92]:

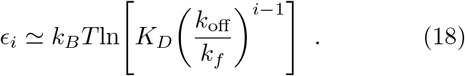

Physically, this expression reflects the fact that the energy of the *i*-th state is the sum of the binding energy of the ligand to the first subunit, |*ϵ*_1_| ≃ ln(*K*_*D*_) and the energies of binding of the subsequent subunits |*ϵ*| ≃ ln(*k*_off_*/k*_*f*_).

Repeating the same process for the KPR receptor with the corresponding conservation constraint 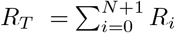 yields the relation

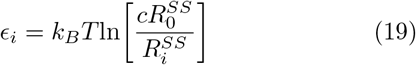

Where 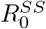 and 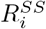 are given by the solution to Eq. 11 at steady state. Ligand binding to all receptor states is given at equilibrium by *k*_on_*c*[*R*_0_] = *k*_off_[*R*_*i*_] so that *ϵ*_*i*_ = *k*_*B*_*T* ln(*K*_*D*_) where *K*_*D*_ = *k*_off_*/k*_on_. This is the mathematical expression of the fact that the equilibrium energetic profile of the KPR receptor is flat.

### C. Variance of the multimer receptor occupancy distribution

We obtain moments of the distribution over multimer receptor states, *P* (***θ***), using derivatives of the free energy. We first assume that the probability of occupancy in state *i* is well approximated by a Gaussian as *P* (***θ***) ∝ *e*^*βF* (***θ***)^. The Cramér-Rao bound gives that var(*x*) ≥ 1*/I*(*x*) where 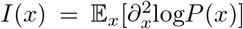 [69, 79]. We use this bound to obtain expressions for the receptor occupancy variance:

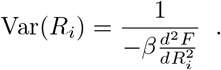

For the signaling complex, *R*_*N*+1_, we use the conservation law 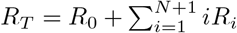 to compute the full derivative 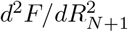 using the partial derivatives:

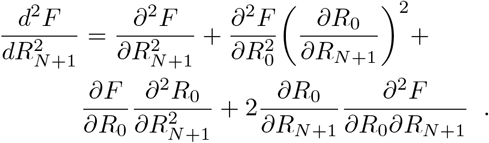

Since 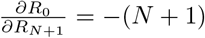 from the conservation law and 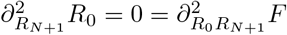, the variance of the multimer receptor states are given, in the Gaussian approximation, by:

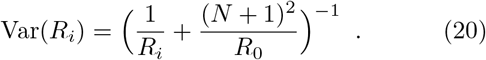

To obtain the variance of the receptor output distribution for the multimeric receptor model we assume that the equilibration time is much shorter than the timescale of cellular decision making. The number of active signaling complexes, *R*_*N*+1_, is modeled as a Gaussian-distributed random variable. This is valid for *R*_*T*_≫ 1 by the Central Limit Theorem, and in practical terms our equations hold well for a few tens of binding events on the cell surface, which is a physiologically realistic assumption [69]. Each signaling complex produces a Poisson-distributed random number of output molecules, *n*. The total signal is then the sum of these *R*_*N*+1_ i.i.d. random variables:

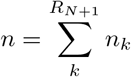

where *k* enumerates the signaling complexes and *n*_*k*_ is the number of molecules produced downstream of receptor *k*. Wald’s equation gives us that the expected total signal is [95, 96]:

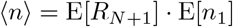

where the expectation is taken for *n*_1_ because all *n*_*k*_ are independent and identically distributed. Since the Poisson process generating output molecules has rate *k*_*p*_*t*, the mean signal has the form

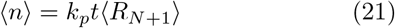

where ⟨*R*_*N*+1_⟩ is the mean number of signaling complexes, obtained from Eqs. 12. The variance in the signal can be obtained using the Blackwell-Girshick equation [96, 97], which states that the variance in the random sum described above is given by

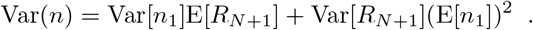

Since the variance of a Poisson process is the same as its mean this yields an expression for the variance of the receptor output:

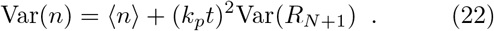

The mathematical expressions for the variance obtained above rely on several simplifying assumptions. The validity of these expressions were tested by comparing predicted means and variances to those obtained from Gillespie simulations of the same model systems [98]. The derived expressions are in good agreement (see Fig. S1).

