## Supplementary Information for "Ligand induced receptor multimerization achieves the specificity enhancement of kinetic proofreading without associated costs"

### I. SUPPLEMENTARY INFORMATION

#### A. Additional details of mathematical methods

The set of ordinary differential equations (ODEs) describing the average dynamics of the multimeric receptor model is:

$$\begin{aligned}
\frac{d[R_0]}{dt} &= -k_{\rm on}c[R_0] + k_{\rm off}([R_T] - [R_0]) \\
\frac{d[R_1]}{dt} &= k_{\rm on}c[R_0] - k_f[R_1] + k_{\rm off}([R_2] - [R_1]) \\
\frac{d[R_2]}{dt} &= k_f([R_1] - [R_2]) - k_{\rm off}[R_2] \\
&\vdots \\
\frac{d[R_{N+1}]}{dt} &= k_f([R_N] - [R_{N+1}]) - k_{\rm off}[R_{N+1}]
\end{aligned} \tag{1}$$

where $[R_0]$ is the in-membrane concentration of unbound receptors, $[R_i]$ for $i > 0$ is the concentration of species formed of $i$ receptors bound in complex by a ligand, $R_T$ is the total number of receptor subunits and is conserved, $\theta$ is set to 1 here but in general can modify dissociation rate constants for in-membrane dissociation reactions, and all rate constants are analogous to those of the KPR model (see also Figure 1 and Section II.A). Analagous to Eq. 3 of the main text, the transition rates for the heteromultimeric receptor are chosen to be:

$$\begin{aligned}
k_{0i} &= k_{\rm on}c \\
k_{i0} &= k_{\rm off} \\
k_{ij} &= e^{\delta_{ij}/k_BT}k_f[R_0] \text{ for } i,j > 0 \\
k_{ji} &= \theta k_{\rm off} \text{ for } i,j > 0
\end{aligned} \tag{2}$$

where $k_{0i}$ and $k_{i0}$ are the association and dissociation rate constants for ligands in the extracellular volume binding/unbinding a single receptor of type $i$, $k_{ij}$ and $k_{ji}$ are the in-membrane association and dissociation rate constants, and $\theta$ allows the in-membrane receptor dissociation to differ from the three dimensional ligand-receptor dissociation rate.

The equilibrium number of heterodimeric ($N = 2$) signaling complexes, used in the main text, is given by:

$$\langle \text{signaling complexes} \rangle = \frac{R_T}{2}\left(1 + \frac{K_4}{R_T}\frac{(c+K_2)(c+K_1)}{cK_1} - \sqrt{(1 + \frac{K_4}{R_T}\frac{(c+K_2)(c+K_1)}{cK_1})^2 - 1 + (\frac{\Delta}{R_T})^2}\right)$$

where $K_1 = k_{\rm off,1}/k_{\rm on}$, $K_2 = k_{\rm off,2}/k_{\rm on}$, $K_4 = \theta k_{\rm off,1}/k_f$, $R_T = R_{1,\rm total} + R_{2,\rm total}$, and $\Delta = R_{1,\rm total} - R_{2,\rm total}$. The mean output from the heterodimeric receptor is $\langle n \rangle = k_p t \langle \text{signaling complexes} \rangle$. The maximum output occurs when $c = \sqrt{K_1K_2}$.

Setting $d[R_i]/dt = 0$ at equilibrium in Supplementary Eqs. 1, the mean number of active signaling complexes can be obtained by solving the system of algebraic equations (Eqs. [12] in the main text).

Prototypical values for biological transition rates for the multimeric receptor are provided in Table S1. Using these values, a typical multimeric receptor may be expected to operate in the range of $\tilde{g} = k_{\rm off}/(k_f[R_T]) \approx 1$ to 10.

| $k_{\rm on}$ | $k_{\rm off}$ | $k_f$ (multimer) | $k_f$ (KPR) | $R_T$ | A |
|---|---|---|---|---|---|
| $10^5 M^{-1}s^{-1}$ | $1s^{-1}$ | $1\mu m^2$molecules$^{-1}s^{-1}$ | 0.36 $s^{-1}$ | 100 to 1000 molecules per cell | 1000 $\mu m^2$ |

TABLE S1. Prototypical values for transition rates, taken from [*Kirby et al. Frontiers. (2021)*, *Pettmann et al. (2021)*]. The number of receptors on a cell can vary by at least an order of magnitude either way depending on the particular cell type and receptor in question.

The dose-response curve of the heterodimeric receptor and an analogous homodimeric receptor are shown in Figure S2. The dose-response curve for the heterodimeric receptor is much broader than the homodimeric receptor, as discussed in the main text.

The mean signaling dynamics of the adaptive sorting model were obtained by numerical integration of the following system of equations:

$$\begin{aligned}
\frac{d[E]}{dt} &= \alpha([C_1]+[D_1])([S_T]-[E]) - \beta[E] \\
\frac{d[C_0]}{dt} &= k_{\text{on}}([L_1]-\sum_{i=0}^{N}[C_i])([R_T]-\sum_{i=0}^{N}[C_i]+[D_i])+ \\
&\quad (b+\gamma[E])[C_1]-(k_f+k_{\text{off},1})[C_0] \\
\frac{d[C_i]}{dt} &= k_f[C_{i-1}]+(b+\gamma[E])[C_{i+1}]- \\
&\quad (k_f+b+\gamma[E]+k_{\text{off},1})[C_i] \text{ for } i\in\{1,N-1\} \\
\frac{d[C_N]}{dt} &= k_f[C_{N-1}]-(b+\gamma[E]+k_{\text{off},1})[C_N] \\
\frac{d[D_0]}{dt} &= k_{\text{on}}([L_2]-\sum_{i=0}^{N}[D_i])([R_T]-\sum_{i=0}^{N}[C_i]+[D_i])+ \\
&\quad (b+\gamma[E])[D_1]-(k_f+k_{\text{off},2})[C_0] \\
\frac{d[D_i]}{dt} &= k_f[D_{i-1}]+(b+\gamma[E])[D_{i+1}]- \\
&\quad (k_f+b+\gamma[E]+k_{\text{off},2})[D_i] \text{ for } i\in\{1,N-1\} \\
\frac{d[D_N]}{dt} &= k_f[D_{N-1}]-(b+\gamma[E]+k_{\text{off},2})[D_N]
\end{aligned}$$

where $[E]$ is the intracellular concentration of enzyme catalyzing the proofreading reaction, $[C_i]$ and $[D_i]$ are the in-membrane concentrations of receptors that have reached the $i^{th}$ state of the proofreading chain while bound to the first or second ligand respectively, $\alpha$ is the rate of state switching for enzyme $[E]$, $b$ is the reverse proofreading rate, $\gamma$ is the equilibrium constant for enzyme $[E]$ binding to the $[C_{N-1}]$ complex, and we used the same parameter values as in [*François and Altan-Bonnet, 2016*] with $N = 5$. The mean signal is then taken to be $k_p t([C_5]+[D_5])$.

A Mathematica notebook with additional computational details is available at: `https://github.com/dakirby/Multimer_Specificity`.

##### B. Mean receptor output from the reversible KPR model

The mean output produced by the reversible KPR receptor is

$$\langle n \rangle = \frac{\nu c e^{\delta_{21}}(k_f + e^{\delta_{02}}k_f + k_{\text{off}}r_0)}{(k_f + e^{\delta_{21}}k_f + e^{\delta_{21}}k_{\text{off}}r_0)(\nu c + e^{\delta_{02}}(k_{\text{off}}+\nu c))}$$

The expression for the variance is impractical to print on a page but a Mathematica notebook with the derivation and full expression is provided at `https://github.com/dakirby/Multimer_Specificity`.

The selectivity metric, $\eta$, for the reversible KPR model in the case $N = 1$ is given by:

$$\eta = \frac{\left(1+e^{\delta_{21}/k_BT}(1+g)\right)\left(x+e^{\delta_{02}/k_BT}(1+x)\right)\left(1+e^{\delta_{02}/k_BT}+g\zeta\right)}{\left(1+e^{\delta_{02}/k_BT}+g\right)\left(x(1+e^{\delta_{02}/k_BT})+e^{\delta_{02}/k_BT}\zeta\right)\left(1+e^{\delta_{21}/k_BT}(1+g\zeta)\right)}$$

where $g = k_{\text{off}}/k_f$, $x = k_{on}c/k_{\text{off}}$, and $k_{on} = r_0\nu$. For $\delta_{02} \gg 1$ and $\delta_{21} \gg 1$ the selectivity scales as $\eta \propto \mathcal{O}(\zeta^{-2})$. The selectivity as a function of proofreading strength, $g$, and relative binding strength, $\zeta$, are shown in Figure S2 (A-B) for the reversible KPR receptor with $N = 1$ and with the homodimeric receptor also shown for comparison.

When $\delta_{02} = -\delta_{21}$ then the Kolmogorov criterion for reversibility is satisfied. In this case,

$$\eta = \left(1+x(1+e^{-\delta})\right)/\left(\zeta+x(1+e^{-\delta})\right)$$

where $\delta$ is now the energy input to suppress both transitions. The selectivity scales as $\mathcal{O}(\zeta^{-1})$ which is the same as a non-proofread receptor. The selectivity, $\eta_1$, as a function of non-equilibrium energetic cost $\delta_{02}$ and (equilibrium) proofreading strength is demonstrated in Figure S2 (C). The non-equilibrium energy only benefits the selectivity enhancement in the regime $k_{\text{off}} \gg k_f$.

##### C. Absolute discrimination with adaptive sorting

One receptor signaling mechanism that has been studied in the context of both ligand discrimination and antagonism is adaptive sorting, an extension of the classical KPR model which has been important for explaining specificity, speed, and sensitivity in TCR signaling [*Lalanne and François, PRL, (2013); François and Altan-Bonnet, J. of Stat. Phys. (2016)*].We introduce adaptive sorting in this section as a point of comparison with multimeric receptors regarding absolute discrimination and antagonism; the speed and sensitivity of adaptive sorting, which are important for the TCR context specifically, go beyond our study of multimeric receptors. Adaptive sorting introduces inter-receptor coupling through an intracellular enzyme that modulates the forward proofreading rate as $k_{ij} = k_f[E]$ where $[E]$ is the enzyme concentration. The concentration of active enzyme is itself affected by the number of receptors bound to either ligand, thereby coupling the response of individual KPR receptors. For a full explanation of adaptive sorting, see [*François and Altan-Bonnet, J. of Stat. Phys. (2016)*].

##### D. Non-equilibrium dimeric receptor model

In Section D of the main text, we consider a non-equilibrium dimeric receptor model. We choose to drive the disassembly of the dimeric receptor complex by a factor $e^{\delta/k_BT}$ because this has the effect of attenuating the signaling state by the same factor. Since KPR works by attenuation of the signaling state, this makes for a functionally comparable effect of $\delta$ in each model. Other choices are certainly possible and physically interesting in their own right. A complete analysis of all possible non-equilibrium drivings lays outside the scope of this paper.

##### E. Specificity can be regulated via non-equilibrium driving

KPR receptors must be driven out of equilibrium to produce specificity enhancement. By contrast, multimeric receptors deliver specificity enhancement already at equilibrium. Yet, non-equilibrium driving may also increase the specificity of the multimeric receptor. To investigate how the specificity and robustness to noise of either mechanism are affected by non-equilibrium energy input, we have generalized the multimer model to include non-equilibrium driving. The non-equilibrium energy is introduced into the multimer model in a manner analogous to KPR: out-of-equilibrum energy input $\delta_{ij}$ drives the receptor away from the signaling state as described in Section IIA of the main text.

As shown in Figure S5, driving either receptor type out of equilibrium increases the specificity. For the KPR receptor, the out-of-equilibrium energy input $\delta > 0$ is necessary to suppress "short-circuiting" transition rates from the unbound state (e.g., $k_{02}$). By contrast, no "short-circuiting" is possible in the multimeric receptor model because all interactions are bi-molecular (there are no loops in the receptor assembly kinetic scheme). The multimeric receptor can thus enhance specificity at equilibrium, although the enhancement is only maximized for $k_{\text{off}}/([R_T]k_f) = \tilde{g} \gg 1$. Figure S5 demonstrates how non-equilibrium energy input can be used to attenuate the multimeric signaling state, effectively compensating for a low equilibrium factor of $\tilde{g}$ in the multimeric model. Naturally, not all non-equilibrium driving increases specificity. In particular, large negative $\delta$ decreases the specificity below the equilibrium specificity for both receptor models because the signaling state is more likely to be occupied regardless of ligand binding strength (see also Figure S1C).

For both models, enhancing specificity by expending energy to decrease the occupancy of the receptor signaling state comes at the cost of a lower SNR. For the dimeric receptor, the SNR scales as $e^{-\delta/2}$ (see Figure S1C) so that higher specificity is achieved at the cost of increased output noise. For the KPR receptor, both $g$ and $\delta$ can attenuate the signaling state and thereby enhance specificity for the receptor. However, altering $g$ is only effective for $\delta > 0$ (see also Figure S2 C).

##### F. Antagonism in ligand mixtures

The predicted signal for dimeric receptors with mixtures of ligands (at concentrations $L_1$ and $L_2$ respectively) is modeled by numerically integrating the system of equations:

$$\frac{d[L_{1,\text{free}}]}{dt} = [L_1] - ([C_0] + [C_1])$$
$$\frac{d[L_{2,\text{free}}]}{dt} = [L_2] - ([D_0] + [D_1])$$
$$\frac{d[R_{\text{free}}]}{dt} = [R_T] - ([C_0] + 2[C_1] + [D_0] + 2[D_1])$$
$$\frac{d[C_0]}{dt} = k_{\text{on}}[L_{1,\text{free}}][R_{\text{free}}] - ([R_{\text{free}}]k_f + k_{\text{off},1})[C_0]$$
$$\frac{d[C_1]}{dt} = k_f[R_{\text{free}}][C_0] - k_{\text{off},1}[C_1]$$
$$\frac{d[D_0]}{dt} = k_{\text{on}}[L_{2,\text{free}}][R_{\text{free}}] - ([R_{\text{free}}]k_f + k_{\text{off},2})[D_0]$$
$$\frac{d[D_1]}{dt} = k_f[R_{\text{free}}][D_0] - k_{\text{off},2}[D_1]$$

using $R_T = 100$ and the other parameter values chosen to match the adaptive sorting model (see main text).

For the KPR receptor in a ligand mixture, steady state equations can be obtained from the system:

$$0 = -(k_{\text{on}}c_1 + k_{\text{on}}c_2)[R_0] + k_{\text{off},1}([R_{1,C}] + [R_{2,C}]) + k_{\text{off},2}([R_{1,D}] + [R_{2,D}])$$
$$0 = k_{\text{on}}c_1[R_0] - (k_{\text{off},1} + k_f)[R_{1,C}]$$
$$0 = k_f[R_{1,C}] - k_{\text{off},1}[R_{2,C}]$$
$$0 = k_{\text{on}}c_2[R_0] - (k_{\text{off},2} + k_f)[R_{1,D}]$$
$$0 = k_f[R_{1,D}] - k_{\text{off},2}[R_{2,D}]$$

subject to the constraint $R_0 + R_{1,C} + R_{2,C} + R_{1,D} + R_{2,D} = R_T$, where $[R_{i,\alpha}]$ denotes the concentration of receptors in state $i$ and bound by ligand $\alpha \in \{C, D\}$, and $c_j$ and $k_{\text{off},j}$ are the corresponding concentration and unbinding rate for ligand $j$ ($j = 1$ for ligand $C$ and $j = 2$ for ligand $D$).

Figure S4 shows that the total signaling state occupancy for both receptor models in ligand mixtures is reduced in the presence of a lower affinity antagonist ligand relative to the same conditions without any antagonist. For both models at low agonist concentrations, the signaling state occupancy starts to recover as the antagonist affinity approaches that of the agonist ($\zeta$ approaches 1). At low agonist concentrations, competitive inhibition is less of an effect and the response is more like the additive response to each ligand. For the KPR receptor, a sufficiently high agonist concentration can overcome the antagonist and saturate the signaling state. For the dimeric receptor, a large agonist concentration cannot overcome the presence of an antagonist because the nature of all multimeric receptor dose-response curves is to decrease at very high ligand concentrations due to self-competition for free receptor subunits.

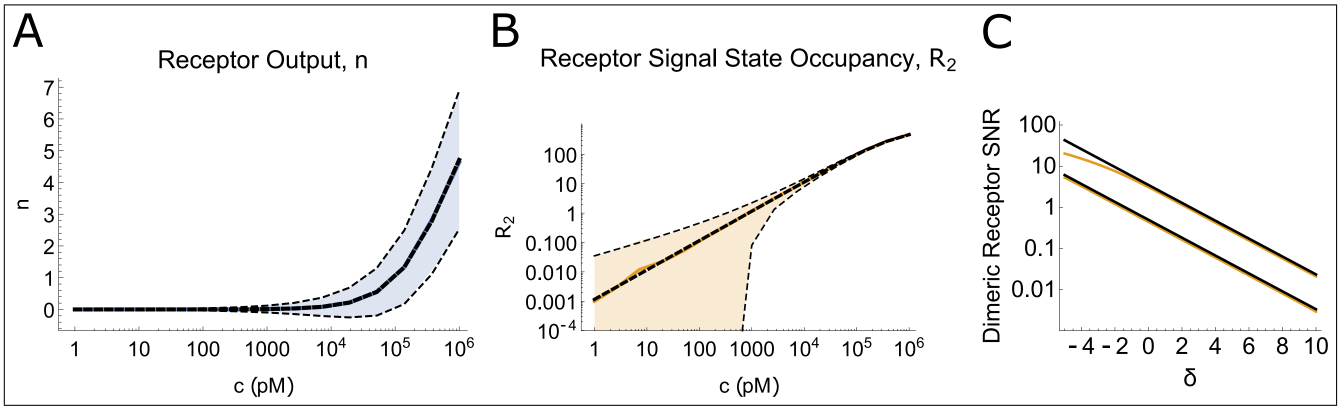


FIG. S1. Validation of multimeric mean and standard deviation expressions. The mean receptor output (A) and steady state number of activated homodimeric receptor complexes (B) as computed from Gillespie simulation are shown as solid lines with the mean ± one standard deviation envelope shaded in colour. The theoretical predictions for the mean of both the output and the signaling complexes are shown as thick dashed lines (overlaid on simulated results), and the predicted one standard deviation envelope is denoted with thin dashed lines. Simulated results and theoretical predictions are in good agreement. Results are from 1000 Gillespie trials. Parameters (taken from [*Kirby et al. Front. (2021)*]) are: $R_T = 4000$ molecules, $A = 2760\mu^2$, $k_p t = 10^2$, $k_{\rm off}/k_{\rm on} = 5\mu M$, $k_{\rm off}/(k_f A) = 1$. C) The theoretical prediction for the non-equilibrium dimeric receptor SNR is plotted (orange lines) for two different conditions: upper line is $k_f[R_T]/k_p = 100$, lower line is $k_f[R_T]/k_p = 1$. The SNR scaling is compared to $\alpha e^{-\delta/2}$ (black lines) where $\alpha$ is a fitted parameter. Other parameters: $k_p t = 100$, $k_{\rm on} c = 10^{-3}$, $k_{\rm off}/k_p = 1$.

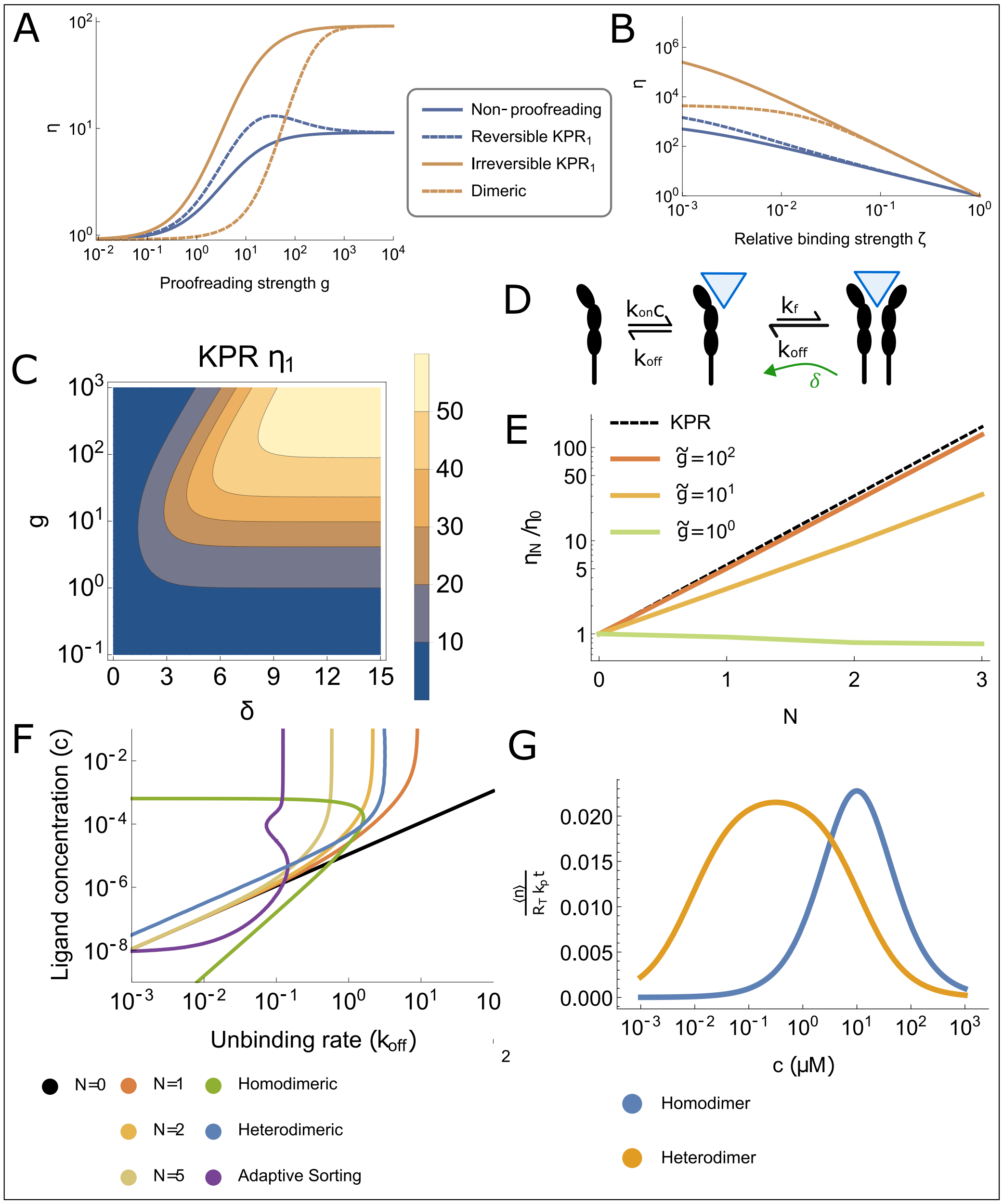


FIG. S2. Additional specificity enhancement plots. A-B) In the small $\delta_{02}$ regime, the specificity of the reversible KPR receptor saturates to the same level as a non-proofreading receptor (dashed blue line), barely enhancing specificity for most proofreading strengths $g$ (A), or differences in ligand affinities $\zeta$ (B). Parameters are $\delta_{02} = 2k_BT$, $\delta_{21} = 0$, $\zeta = 0.1$ for (A), $\frac{k_{on}c}{k_{off}} = 10^{-3}$, $k_{off}/k_f = 10^3$ for (B), and $R_T = 100$. C) The sensitivity $\eta$ for a KPR receptor with $N = 1$ as a function of the proofreading strength $g = k_{off}/k_f$ (fixing $k_{off}/k_p = 1$ and varying $k_f$) and non-equilibrium energy input $\delta$; $k_{on}c/k_{off} = 10^{-3}$ and $\zeta = 0.1$. D) Schematic for the non-equilibrium dimeric receptor model, indicating the effect of non-equilibrium energy input $\delta$ to drive the system away from the signaling state. E) The specificity of the multimeric receptor as $R_T$ changes. Here we show that for $R_T = 10^4$ copies per cell (i.e., $\tilde{g} = 1$) that the specificity enhancement mechanism breaks down. Parameters are otherwise the same as Figure 2B of the main text. F) Ligand concentration $c$ (in units of $k_p/k_{on}$) required to achieve a desired response – fixed at $\langle n \rangle / R_T = 30$ – as a function of the ligand unbinding rate $k_{off}$ (in units of $k_p$). Same as in Figure 4B but now including the adaptive sorting model. For the adaptive sorting model we used parameters from [*Lalanne and François, PRL (2013)*]. G) Heterodimeric receptor dose response curve versus an analogous homodimeric receptor dose response curve. Parameters taken from Table S1 and for the heterodimeric receptor $k_{off,1} = 1s^{-1}$, $k_{off,2} = 10^{-3}s^{-1}$ which represents the physiological range of Type I Interferon receptor affinities.

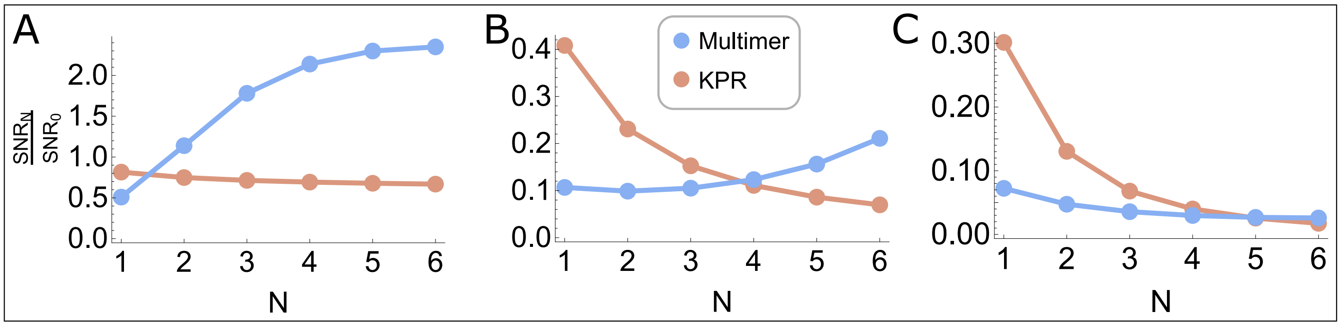


FIG. S3. The SNR of each receptor model relative to that of the $N = 0$ model, as a function of $N$. Parameters the same as Fig. 5 of the main text except $k_{\rm off}$ chosen to give: A): $\tilde{g} = g = 1/(N+1)$. B): $\tilde{g} = g = 10/(N+1)$. C): $\tilde{g} = g = 20/(N+1)$.

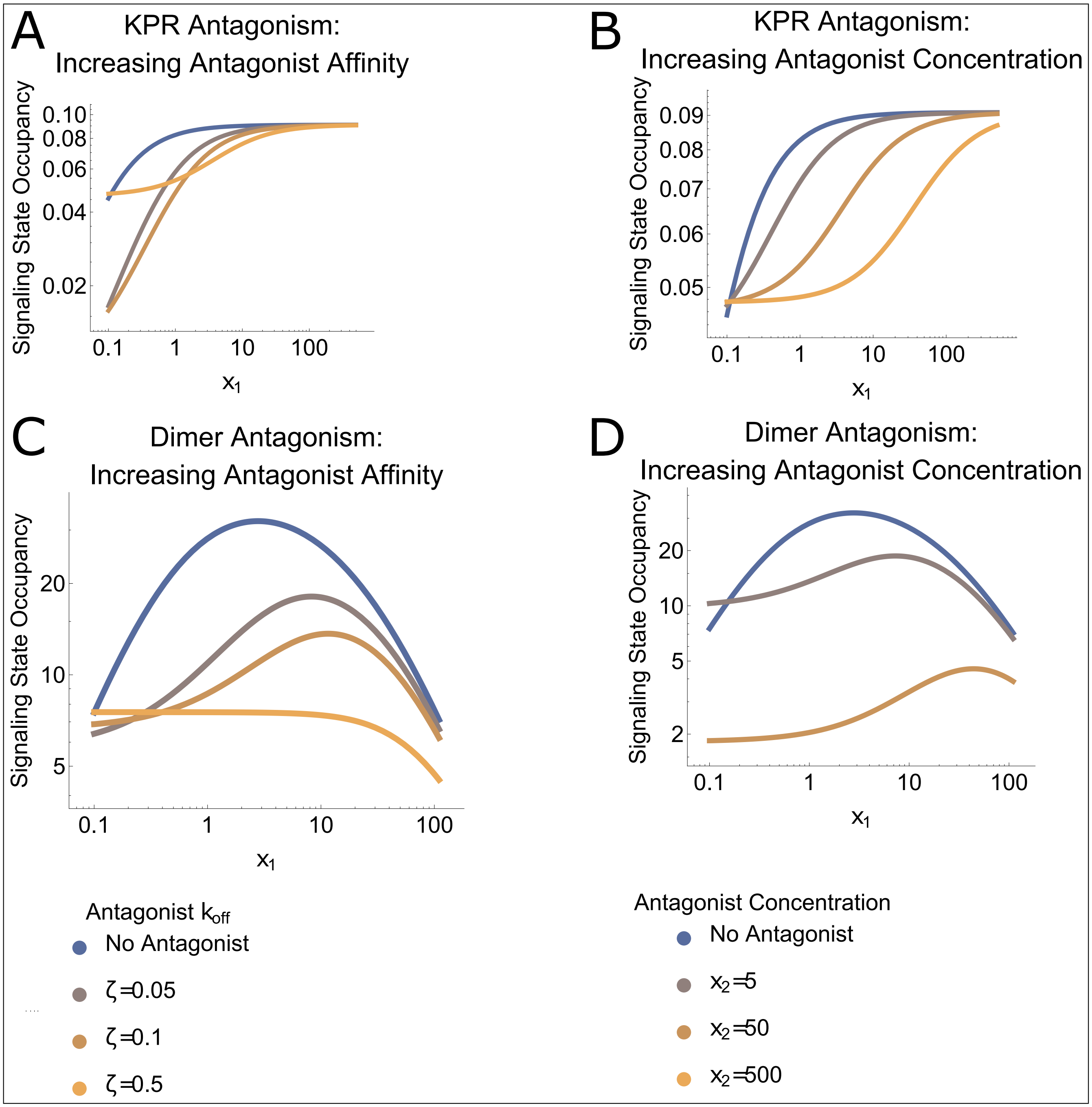


FIG. S4. Dimeric and 1-step KPR receptors exhibit antagonism. The signaling state occupancy, $R_2$, as a function of agonist concentration $x_1 = k_{\rm on}c_1/k_{\rm off,1}$. (A) KPR response for increasingly strong binding antagonist. (B) KPR response for increasing antagonist concentration. (C) Dimeric receptor response for increasingly strong binding antagonist. (D) Dimeric receptor response for increasing antagonist concentration. Antagonist concentration $x_2 = 500$ is omitted because the occupancy is so low. For (A) and (C): $k_{\rm on}/k_{\rm off,1} = 10$, $k_{\rm off}/k_f = 10$, $[R_T] = 100$,and $k_{\rm on}c_2 = 10$. For (B) and (D): same as above but $c_2$ varies and $k_{\rm on}/k_{\rm off,2} = 5$.

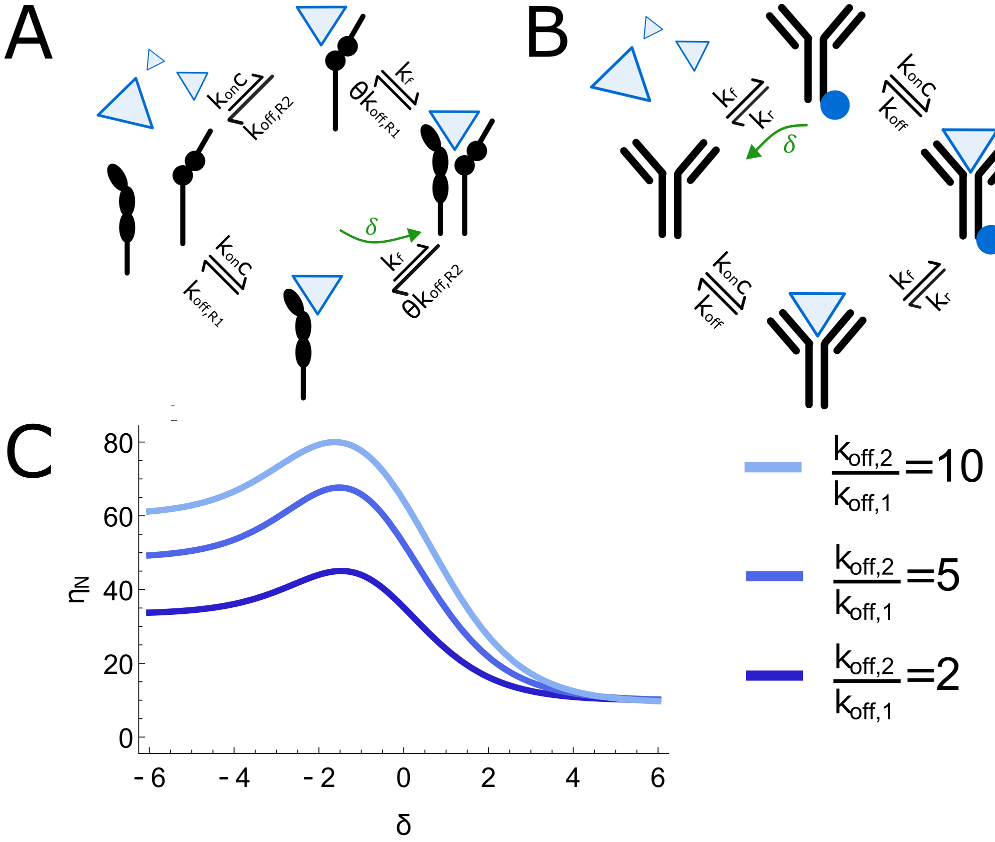


FIG. S5. Non-equilibrium driving modulates the signaling performance. A) The heterodimeric receptor, repeated from Fig. 3A for convenience. B) The non-equilibrium KPR receptor may be modeled in a similar fashion to the heterodimeric receptor as a four state-model with significant non-equilibrium driving to inhibit both the reactions taking the receptor from bound states $i+1$ to $i$ and the reactions taking the receptor from the empty state to the signaling state (i.e., state 0 to state $N+1$). For large enough $\delta$, this model has identical signaling behaviour to the KPR model considered in the main text. C) The non-equilibrium driving $\delta_{ji}$ can accentuate the specificity enhancement by a heterodimeric receptor. In this case, the driving alters the rate of reaction for $R_2$ association with the ligand-$R_1$ complex. The specificity enhancement is strongest when $k_{\text{off},2} \gg k_{\text{off},1}$ and when the association is inhibited ($\delta < 0$) but not so strong as to prevent the formation of the signaling complex altogether. Parameters the same as Fig. 3C except $k_{\text{off},2}$ is varied.
